# Differential antibody dynamics to SARS-CoV-2 infection and vaccination

**DOI:** 10.1101/2021.09.09.459504

**Authors:** Yuezhou Chen, Pei Tong, Noah B. Whiteman, Ali Sanjari Moghaddam, Adam Zuiani, Shaghayegh Habibi, Avneesh Gautam, Tianshu Xiao, Yongfei Cai, Bing Chen, Duane R. Wesemann

## Abstract

Optimal immune responses furnish long-lasting (durable) antibodies protective across dynamically mutating viral variants (broad). To assess robustness of mRNA vaccine-induced immunity, we compared antibody durability and breadth after SARS-CoV-2 infection and vaccination. While vaccination delivered robust initial virus-specific antibodies with some cross-variant coverage, pre-variant SARS-CoV-2 infection-induced antibodies, while modest in magnitude, showed highly stable long-term antibody dynamics. Vaccination after infection induced maximal antibody magnitudes with enhanced longitudinal stability while infection-naïve vaccinee antibodies fell with time to post-infection-alone levels. The composition of antibody neutralizing activity to variant relative to original virus also differed between groups, with infection-induced antibodies demonstrating greater relative breadth. Differential antibody durability trajectories favored COVID-19-recovered subjects with dual memory B cell features of greater early antibody somatic mutation and cross-coronavirus reactivity. By illuminating an infection-mediated antibody breadth advantage and an anti-SARS-CoV-2 antibody durability-enhancing function conferred by recalled immunity, these findings may serve as guides for ongoing vaccine strategy improvement.

## INTRODUCTION

The devastating effects of the COVID-19 pandemic can be attenuated by mRNA vaccines encoding for SARS-CoV-2 spike (S) (Krause et al., 2021; Thompson et al., 2021). However, emergence of mutated variants (Hacisuleyman et al., 2021) and waning immunity (Pegu et al., 2021; Shrotri et al., 2021) are continuing major threats. Strategies to maximize robust levels of protective antibodies that are stable over time and retain function across mutating variants through vaccination is a vital goal. Understanding the capabilities of the immune system in this regard, and how available vaccines can elicit them is a critical step in this process, both for the ongoing SARS-CoV-2 pandemic as well as for vaccine strategies more broadly.

Antibodies are B cell-expressed molecules composed of immunoglobulin (Ig) heavy (H) and light (L) chains and are produced in the context of different IgH isotypes (e.g. IgM, IgG, IgA). Induced through infection or vaccination, antibodies can evolve to better recognize the pathogen through B cell clonal selection and somatic hypermutation (SHM), and subsequently produce affinity-matured versions of antibodies at greater speed and magnitude, which provides enhanced immunity against similar future threats. After infection or vaccination, IgG antibodies can be sustained for decades (Amanna et al., 2007). Durable antibody responses require coordinated T and B cell interactions within germinal centers (GCs) (Cyster and Allen, 2019; Mesin et al., 2016). GC B cells can differentiate into long-lived plasma cells (LLPCs) or memory B cells. Memory B cells can more efficiently differentiate into antibody secreting plasma cells upon subsequent pathogen invasion, but pre-formed pathogen-specific antibodies produced from LLPCs are prophylactically available before repeat pathogen invasion attempts, and thus provide immediate protection. B cells that are activated outside of GCs can also differentiate into memory B cells (Takemori et al., 2014) in addition to shorter-lived versions of antibody-secreting cells.

SARS-CoV-2 infection and vaccination elicits neutralizing antibodies targeting the viral spike glycoprotein (S), a homotrimer of precursor polypeptide chains that are cleaved upon maturation into two fragments, S1 and S2. The S1 region contains the receptor binding domain (RBD) and the N-terminal domain (NTD), both targets of antibodies with potent neutralizing capability. The S2 region is generally conserved across coronaviruses and mediates viral membrane fusion required for entry to host cells (Abbott and Crotty, 2020; Hoffmann et al., 2020; Letko et al., 2020; Walls et al., 2020; Wrapp et al., 2020).

Anti-S neutralizing antibodies are key correlates of disease protection (Addetia et al., 2020; Khoury et al., 2021; Lopez Bernal et al., 2021; Shrotri et al., 2021). In this context, declining antibodies over time and immune escape have driven the discussion on needs for evolution of vaccine strategies, such as additional dosing (Bergwerk et al., 2021; Garcia-Beltran et al., 2021; Hoffmann et al., 2021; Lopez Bernal et al., 2021; Planas et al., 2021; Shrotri et al., 2021; Zhou et al., 2021). Understanding factors that can maximize anti-SARS-CoV-2 antibody durability and breadth will be critical to inform ways for vaccine strategy optimization. To this end, we charted virus-specific antibody durability and cross-variant breadth and examined how these features dynamically change over time after infection versus mRNA vaccination. We also examined how prior infection influences vaccine responses. We focused our analysis on features of cross-variant antibody breadth as well as signatures that may endow antibody responses with greater antibody durability.

We report here that antibodies induced by natural infection had robust long-term stability at modest levels and greater polyclonal neutralizing breadth across viral variants compared to infection-naïve vaccinees. On the other hand, vaccination alone induced robust initial virus-specific antibodies with some cross-variant coverage, and vaccination after infection induced the greatest antibody magnitudes with enhanced longitudinal stability over time. Infection-naïve vaccinee antibodies declined with time to post-infection-alone levels despite two doses, while those that recovered from early pandemic SARS-CoV-2 infection (i.e. before the emergence of Alpha, Beta, Gamma, Delta (Kupferschmidt, 2021; Summer E. Galloway, January 22, 2021)) elicited antibodies with greater breath across new viral variants than antibodies from infection-naïve vaccinees. Differential antibody durability trajectories favored COVID-19 convalescent subjects with dual memory B cell features of greater antibody somatic mutation and cross-coronavirus reactivity. Collectively, these studies identify an antibody breadth enhancing ability of early pandemic (i.e., pre-variant) SARS-CoV-2 infection, and a durability-enhancing function conferred by recalled immunity.

## RESULTS

### Vaccine recall of prior infection confers greater antibody magnitude and stability

To gain insights into how immune recall influences antibody durability, we charted anti-S and anti-RBD IgG dynamics in three longitudinal cohorts: (i) Individuals who have recovered from SARS-CoV-2 infection that had not yet been vaccinated (COVID-19 convalescents), (ii) COVID-19 convalescents who had received mRNA vaccination (COVID-19 vaccinees), and (iii) COVID-19-naïve individuals that had received mRNA vaccination (naïve vaccinees). COVID-19 convalescents (n=62) were recruited between March 2020 and June 2020 in Boston (Chen et al., 2020), before the emergence of the Alpha, Beta, Gamma, and Delta variants (Kupferschmidt, 2021; Summer E. Galloway, January 22, 2021). Blood was drawn over the course of approximately 6-9 months with 5 or 6 blood draws (Figure S1A and Table S1). Twenty-eight COVID-19 convalescents continued to provide blood donations after mRNA vaccination (Table S2, Figure S1B-G). Eighteen naïve vaccinees donated up to 8 repeated blood draws after mRNA vaccination until ~195 days following the second dose (Table S3, Figure S1B-G).

Anti-S and Anti-RBD IgG in COVID-19 convalescents had the stablest antibody levels over time maintaining most (60-80%) of peak levels by ~220 days after symptom onset (Figure 1A, B). Consistent with previous reports (Planas et al., 2021; Reynolds et al., 2021; Stamatatos et al., 2021), one dose of mRNA vaccination boosted anti-S and anti-RBD IgG in COVID-19 vaccinees to maximal levels, beyond levels attained with two doses from naïve vaccinees (Figure 1A, B).

**Figure 1.**
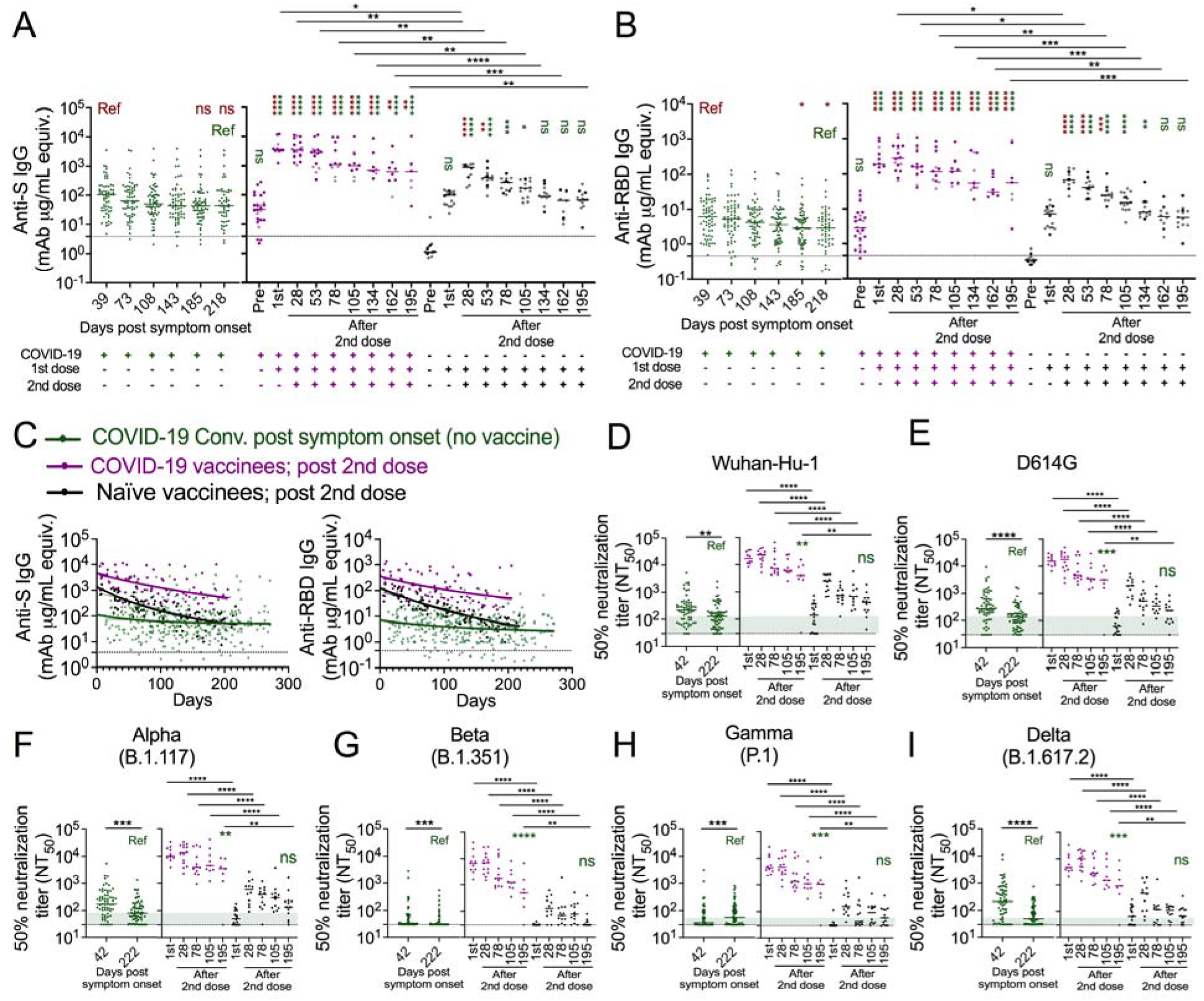
Vaccine recall of prior infection confers greater antibody magnitude and stability. (A, B) Dot plots showing anti-S (A) and anti-RBD (B) IgG antibody levels in seroconverted COVID-19 convalescents (green, n = 62, left), COVID-19 vaccinees (purple, n = 7-28), and naïve vaccinees (black, n = 10-18) over time as indicated. For vaccinees, darker color indicates Moderna mRNA-1273 vaccination and lighter color indicates Pfizer-BioNTech BNT162b2 vaccination. Pre-vaccination plasma (Pre) from COVID-19 convalescents was collected at a median of 222 days (range 189-271) after symptom onset and a median of 76 days (range 13-187) before the 1^st^ vaccine dose (1st). After vaccination, subjects donated 2-8 longitudinal blood draws up to 217 days after the 2^nd^ vaccine dose. Blood collections were performed at a median day 22 (range 10-31) after the 1^st^ vaccination. After second vaccination, blood draws were collected at the indicated median days. (C) Dot and line graphs showing anti-S IgG (left) and anti-RBD IgG (right) trajectories in COVID-19 convalescents after natural infection (green, n = 62), and COVID-19 vaccinees (purple, n =28) as well as naïve vaccinees (black, n = 18) after the 2^nd^ vaccine dose. Overlapping fitted curves for both one phase decay (thick line) and linear regression (thin line) models are shown. Green line indicates one-phase decay model and linear regression model fitted curves for log transformed antibody data from the COVID-19 convalescents (anti-S IgG: R = −0.11, peak level =2.04, plateau = 1.69, slope = - 0.00082; anti-RBD IgG: R = −0.17, peak level = 0.86, plateau = 0.39, slope = −0.0013); Purple line indicates COVID-19 vaccinees (anti-S IgG: R = −0.51, peak level = 3.66, plateau = 1.94, slope = −0.0047; anti-RBD IgG: R =-0.46, peak level = 2.57, plateau = 1.19, slope = −0.0042); black line indicates vaccinated naïve subjects (anti-S IgG: R = −0.79, peak level = 3.14, plateau = 1.27, slope = −0.0065; anti-RBD IgG: R = −0.82, peak level = 2.10, plateau = −0.31, slope = - 0.0067). The fitted curves from the three groups were significantly different with each other (p<0.0001). The Dashed lines represent twice the average of pre-COVID (negative) controls. Extra sum-of squares F test found no significant differences between one phase decay and simple linear regression fitted curves in all cases except anti-S IgG log transformed data from naïve vaccinees. (D-I) Dot plots showing pseudovirus neutralization titers to indicated SARS-CoV-2 variants in plasma collected from COVID-19 convalescents after natural infection (green, n = 62), and COVID-19 vaccinees (purple, n =9-27) as well as naïve vaccinees (black, n = 12-18) after second vaccination as indicated. Red and green asterisks indicate comparisons to early (red) and late (green) COVID-19 convalescent antibody levels as indicated by the red and green “Ref”. Dashed black lines in A-C represent twice the average of negative controls, and dashed black lines in D-I represent the limit of neutralization detection (i.e. 30). The light green shading in panels D-I indicates the difference between the assay detection limit and the median neutralization level in COVID-19 convalescents at median day of 222 days after symptom onset. NOVA with Tukey’s multiple comparison test was performed to test significant difference of log-transformed antibody data. Extra sum of squares F test was used to test for significance of differences between the one phase decay model fitted curves. Analysis of Covariance (ANCOVA) was performed to test the significance of differences between slopes generated from linear regression. All other significance testing used the Mann-Whitney U test. *p<0.05, **p<0.01 ***p<0.0005; ****p<0.0001

Both COVID-19 vaccinees and naïve vaccinees had higher peak anti-S and anti-RBD IgG than did naturally infected subjects early after SARS-CoV-2 infection (~40 days post symptom onset) (Figure 1A, B). We tracked antibody responses in mRNA vaccinees for up to 7 months. Consistent with higher peak antibody level after the 2^nd^ dose, COVID-19 vaccinees had significantly higher anti-S and anti-RBD IgG than naïve vaccinees and naturally infected subjects at all plasma collection points out to 7 months (Figure 1A, B). The level of naïve vaccinee anti-S IgG at ~134 days after second dose decreased to the level found in ~220-day unvaccinated COVID-19 convalescents (Figure 1A, B).

Both linear regression and one phase decay models can fit dynamic antibody data reasonably well (Dan et al., 2021) and were used here to provide insights into peak/plateau characteristics (one phase decay, thick line) and dynamic stability (linear regression, thin line) (Figure 1C).

Both models fit the data well with overlapped fitting lines (Figure 1C). While subjects with natural infection had modest anti-S and anti-RBD IgG peak levels, over time they exhibited a high degree of dynamic stability with significantly more stable linear regression slopes than COVID-19 vaccinees and naïve vaccinees (anti-S IgG: −0.00082, p = 0.0026, p < 0.0001; anti-RBD IgG: −0.0013, p = 0.0125, p < 0.0001) (Figure 1C). In contrast, more striking antibody decay was observed in naïve vaccinees. While anti-S and anti-RBD IgG peaked an order of magnitude higher in naive vaccinees than in naturally infected subjects, the higher levels were not maintained over time, coming down to levels on par with COVID-19 convalescent long-term plateau levels. In contrast, COVID-19 vaccinees had the highest level of peak antibodies among the three groups, as well as a more stable plateau phase over time than naive vaccinees. In general, the Pfizer-BioNTech vaccine induced slightly lower anti-S and anti-RBD IgG than the Moderna vaccine, but the difference did not reach statistical significance (Figure S2 A, B) and the trajectory dynamics were similar (Figure S2 C-F).

To examine functional antibody breadth, we compared cross-variant neutralization in the plasma using pseudotyped virus neutralization assays we have previously shown to mirror that seen with authentic virus (Tong et al., 2021). We found that COVID-19 vaccinee plasma had greater cross-variant neutralization function than did plasma from naïve vaccinees and unvaccinated COVID-19 convalescents at all tested time points (Figure 1D-I), consistent with greater anti-S and anti-RBD IgG levels in this group. Six months after the 2^nd^ dose, neutralization levels to all variants in naïve vaccinees decreased to the level found in naturally infected subjects at ~220 days after symptom onset (Figure 1D-I). Together, these data suggest that mRNA vaccines induced higher peak but not greater long-term anti-S and anti-RBD IgG in naïve vaccinees than in naturally infected individuals. In addition, pre-existing immunity from natural infection primed greater vaccine-induced magnitude and long-term stability.

### Greater cross-variant neutralization breadth induced by natural infection

We next asked how recognition breadth to variants evolved after natural infection and mRNA vaccination. To normalize for antibody magnitude, we generated a breadth index which was the quotient of 50% neutralization titer (NT_50_) to each variant divided by NT_50_ to original Wuhan-Hu-1 strain for each plasma sample (Figure 2A). From ~40 days to ~220 days post symptom onset, the breadth index in plasma collected from subjects after natural infection evolved over time (Figure 2B). Compared to samples collected early after infection, COVID-19 convalescents had significantly greater neutralization breadth to the Gamma variant and lower neutralization breadth to the Delta variant ~220 days after symptom onset (Figure 2B). Neutralization function against the Beta variant was lowest among all the tested variants, suggesting that this variant was most resistant to neutralizing antibodies (Figure 2B).

**Figure 2.**
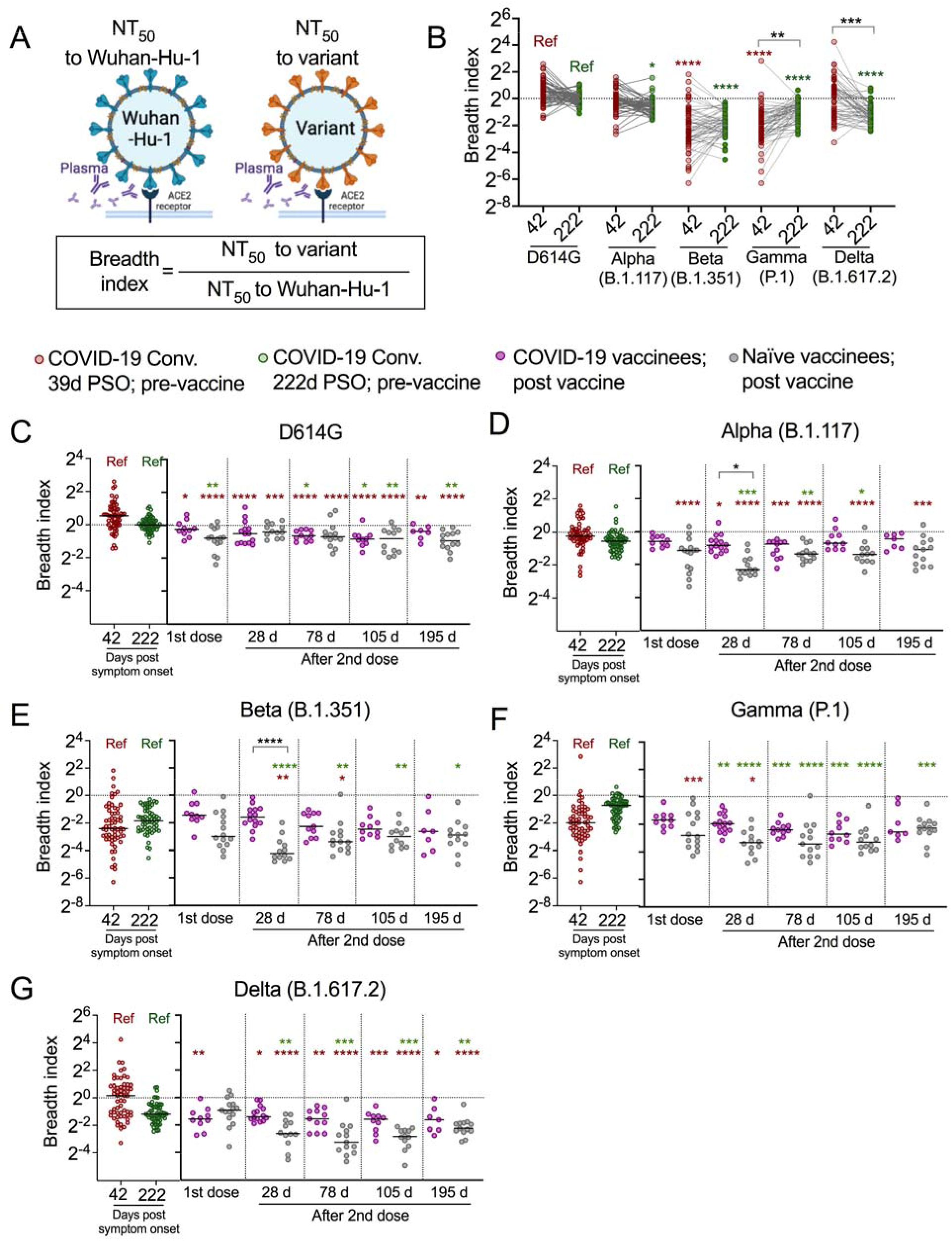
Greater cross-variant neutralization breadth induced by natural infection. (A) Schematics to show the breadth index calculation, which is the ratio of the 50% neutralization titer (NT_50_) to SARS-CoV-2 variant divided by NT_50_ to the original Wuhan-Hu-1 strain of the same plasma sample. (B) Dot and line graphs of calculated breadth indexes to the indicated variants in plasma polyclonal neutralizing antibodies collected at median 42 days (red, n = 59-60) and 222 days (green, n = 57-60) after COVID-19 symptom onset before vaccination. (C-G) Dot plots showing breadth indexes to the indicated SARS-CoV-2 variants in plasma collected from COVID-19 convalescents at the indicated median time points (days) as described in (B), COVID-19 vaccinees (purple, n = 7-14), and naïve vaccinees (grey, n = 12-15). Red and green asterisks indicate comparisons to early (red) and late (green) COVID-19 convalescent breadth indexes as indicated by the red and green “Ref”. Kruskal-Wallis test was performed to test significance in multiple comparisons. *p<0.05, **p<0.01 ***p<0.0005; ****p<0.0001

When comparing COVID-19 convalescents and naïve subjects after vaccination, we found unvaccinated COVID-19 convalescents had significantly greater breadth indexes to all tested variants of concern (VOC) than naïve vaccinees (Figure 2C-G). In addition, vaccination of COVID-19 convalescents did not provide further enhancement of the breadth index for any variant (Figure 2C-G). The differences of breadth indexes to variants between COVID-19 vaccinees and naïve vaccinees were most striking early after the second dose, and gradually decreased afterwards (Figure 2C-G). First dose vaccination in naïve subjects induced similar breadth index values to the Delta variant as naturally infected subjects (Figure 2G), but the second dose significantly decreased the breadth index to this variant in naïve subjects (Figure 2G). Relative to original Wuhan-Hu-1 strain, these results indicate that natural infection elicited a polyclonal antibody response with greater recognition breadth than mRNA vaccination, and that mRNA vaccination with Wuhan-Hu-1 may be unable to enhance breadth beyond that elicited by pre-VOC SARS-CoV-2 infection.

### Early memory B cell SHM and cross-reactivity are signatures of greater antibody durability

Because prior SARS-CoV-2 infection enhanced durability after vaccination, we considered whether prior coronavirus infection may have influenced differential antibody durability in our convalescent cohort. We previously reported two general subsets of COVID-19 convalescents with respect to antibody dynamics over time, antibody sustainers and antibody decayers (Chen et al., 2020), subsequently validated in an independent cohort (Ortega et al., 2021). Sustainers exhibited the same or increasing antibody levels over time while decayers were defined as those that lost antibody levels over the same time frame. Because pre-disease blood draws are not available in the COVID-19 convalescent cohort, we used earliest available blood draws (~40 days post symptom onset) to ask whether there are any differences in other CoV cross reactivity in the early post-infection memory B cell compartment of sustainers versus decayers.

As described previously (Chen et al., 2020), we generated durability indexes by taking virus-specific IgG magnitude values from the most recent blood draw (5^th^ or 6^th^, median 222, range 185-271 days post symptom onset) divided by the respective IgG levels of the 1^st^ draw (median 39, range 13-85 days post symptom onset) for nucleocapsid (N), S and RBD (Figure 3A). Anti-S IgG durability index grouped COVID-19 convalescents into long-term sustainer (durability index ≥1, n = 18) and decayer (durability index < 1, n = 44). Sustainers had shorter symptom duration and milder disease severity as we reported previously (Figure S3 A) (Chen et al., 2020). The sustainers had more anti-S and anti-RBD IgG than decayers, for up to 9-months after symptom onset (Figure 3 B, C).

**Figure 3.**
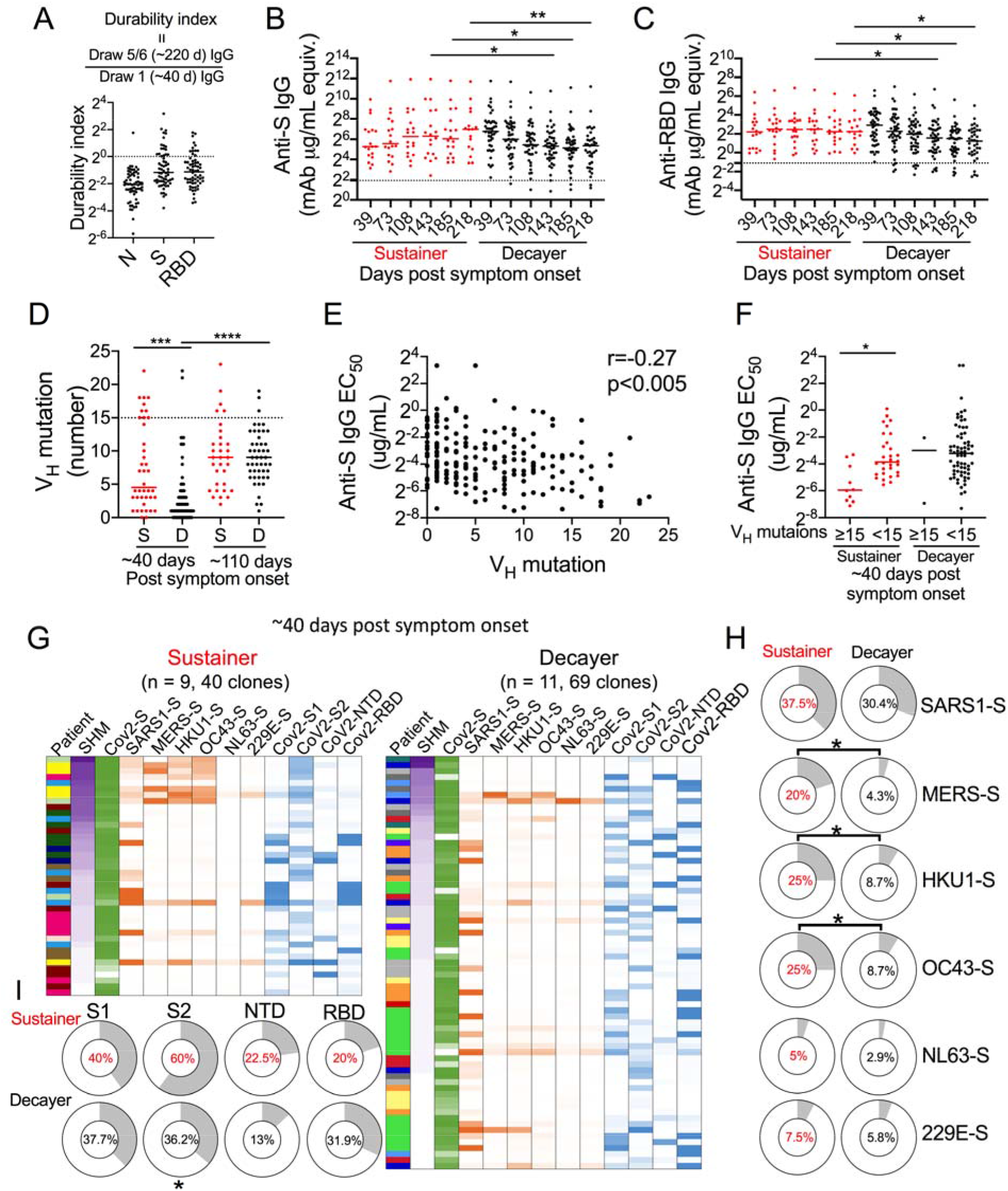
Early memory B cell SHM and cross-reactivity are signatures of greater antibody durability. (A) Dot plot illustrating anti-N, S and RBD antibody durability indexes from seroconverted subjects (n=62). The durability index was reported as the ratio of the last draw (5^th^ or 6^th^ draw) IgG level to the first draw IgG level of same subject. Dashed line represents durability index of 1. (B, C) Dot plots showing anti-S (B) and anti-RBD (C) IgG levels in sustainers (anti-S IgG durability index ≥ 1, red, n = 16-18) and decayers (anti-S IgG durability index <1, red, black, n = 37-44). Dashed lines represent twice the average of pre-COVID controls. Mann-Whitney U test. (D) Dot plot showing mAb heavy chain V gene segment (V_H_) mutation number per sequence cloned from S^+^ memory B cells in sustainer (S, red) and decayer (D, black) collected ~ 40 days after symptom onset (sustainer, n = 9, 40 clones; decayer, n = 11, 69 clones;1^st^ draw) and ~110 days after symptom onset (sustainer, n = 8, 32 clones; decayer, n = 9, 48 clones; 3^rd^ draw). Kruskal-Wallis test. (E) Scatter plots illustrating Spearman correlation between anti-S IgG EC_50_ and V_H_ mutation in all mAbs described in (D). (F) Half-maximal effective concentration of binding (EC_50_) data of sustainer-isolated (red) versus decayer-isolated (black) anti-S mAbs with greater than or less than 15 V_H_ mutations as indicated. The mAbs were isolated ~ 40 days after symptom onset. Kruskal-Wallis test. (G) Heatmap of SHM and binding analysis of mAbs cloned from sustainers (left) and decayers (right) collected ~40 days after symptom onset as described in (D). mAbs are listed in order of highest to lowest SHM. SHM amount is indicated by color (purple) intensity. Different colors in the first column for each panel indicate different individuals. Green color intensity indicates mAb binding avidity to SARS-CoV-2 spike from ELISA analysis. Orange indicates mAb binding to HCoV spikes. Blue indicates mAb binding to SARS-CoV-2 subunits. Intensity of color indicates OD_405_ values. (H) Donut charts showing percentages of mAb binding to the indicated HCoV spikes in sustainer (red) and decayer (black) groups as described in (G). Fisher exact test. (I) Donut plots showing percentages of mAb binding to the indicated SARS-CoV-2 S subdomains in sustainer (red) and decayer (black) groups as described in (G). Fisher exact test. *p<0.05, **p<0.01, ***p<0.005.

T cell crossreactivity analysis revealed that, while sustainers demonstrated higher frequency of non-naïve CD4^+^ (CD45RA^-^CD4^+^) T cells than decayers (Chen et al., 2020), there were no significant differences in antigen specific CD4^+^ T cell or CD8^+^ T cells between sustainers and decayers (Figure S3 B-D) as determined by activation induced marker (AIM) assays (Grifoni et al., 2020; Mateus et al., 2020) (Figure S3 B-D).

To assess antibody cross-reactivity to other coronaviruses, we isolated S^+^ memory B cells, cloned antibody genes, and purified their monoclonal antibodies (mAb) from sustainers and decayers ~40 and ~110 days post symptom onset (Chen et al., 2020). Individuals for memory B cell sorting were selected based on similar initial antibody levels (Chen et al., 2020). We previously showed that by ~110 days, sustainers had higher SHM in their S^+^ memory B cell antibody genes early after infection (~40 days) than decayers (Chen et al., 2020). Subsequent analysis showed that these sustainers retained anti-S or anti-RBD durability indexes at levels ≥1 to at least ~220 days after symptom onset. We produced mAbs from a subset of these clones from which we were able to recover paired heavy and light chain sequences from single cell antibody cloning. This subset showed a higher level of SHM in sustainers at day ~40 (Figure 3D), like the sequence analysis of the overall pool (Chen et al., 2020). When compared to mAbs cloned and produced from samples drawn at ~40 days after symptom onset, those from ~110-day samples had higher SHM as expected (Chen et al., 2020).

We examined the binding of each mAb to spike expressed on 293T cells by flow cytometry and estimated their avidity by calculating half-maximum effective concentration of binding (EC_50_). We found that S^+^ mAb EC_50_s were significantly negatively correlated with IgH V gene mutations, indicating that more highly mutated mAbs had higher avidity for S (Figure 3E). Sustainers had significantly greater number of highly mutated clones with at least 15 IgH mutations ~40 days after symptom onset, as expected (Chen et al., 2020) (Figure 3F).

mAb binding analysis showed that sustainers harbored significantly higher frequencies of memory B cells reactive with spike proteins from seasonal human coronaviruses (HCoVs) HKU1 and OC43, and these tended to have higher SHM and map to S2 (Figure 3G, H). In addition, sustainers had significantly higher frequency of overall memory B cells binding to S2 (Figure 3G, I), which is highly conserved across HCoVs (Guthmiller and Wilson, 2020; Ng et al., 2020; Shrock et al., 2020). No significant differences were observed in the amounts of plasma antibody to HCoV spike between sustainers and decayers (Figure S3E).

At ~110 days after symptom onset, the frequency of S2 and seasonal HCoV cross-reactive clones tended to decrease and the frequency of S1 and RBD binding clones to increase (Figure S4A). The S2/RBD plasma IgG ratio in sustainers early after infection tended to be greater in sustainers than decayers at ~40 days in the total population (Fig. S4B)—a trend that achieved significance in plasma linked to the subset of individuals from which antibodies of S^+^ memory B cells were cloned and expressed (Fig. S4C). The greater S2/RBD IgG in early, but not late sustainer plasma after infection was mirrored in a similar S2/RBD ratio analysis of sustainer and decayer memory B cells (Figure S4D). These findings are consistent with the idea that higher relative proportion of anti-S2 IgG in sustainers early after infection may suppress recalled S2-binding cross-reactive memory B cell frequencies over time, perhaps due to an epitope masking phenomenon by pre-existing anti-S2 antibodies, driving a change in immunodominance over time (Angeletti et al., 2017). Together, these data suggest that early cross-reactive memory B cells recalled from prior CoV infection is linked to antibody dynamic stability over time.

### Anti-S IgG sustainers produced superior cross-variant neutralizing capability

To assess whether the antibody sustainer phenotype has an influence on long-term neutralization capacity, we evaluated sustainer versus decayer cross-variant SARS-CoV-2 neutralization ability of the plasma collected ~40 days (draw 1 or 2, median 42, range 13-119) and ~220 days (draw 5 or 6, median 222, range 185-271) after symptom onset. We calculated NT_50_ for each combination of plasma and SARS-CoV-2 variant pseudovirus (Figure 4 and S4 EG). While there was no NT_50_ difference in plasma collected ~40 days after symptom onset between sustainers and decayers, neutralizing antibody levels to original Wuhan-Hu-1 strain as well as variants of concern Alpha, Beta, and Delta were significantly greater in sustainers compared to decayers at ~220 days after symptom onset (Figure 4A).

**Figure 4.**
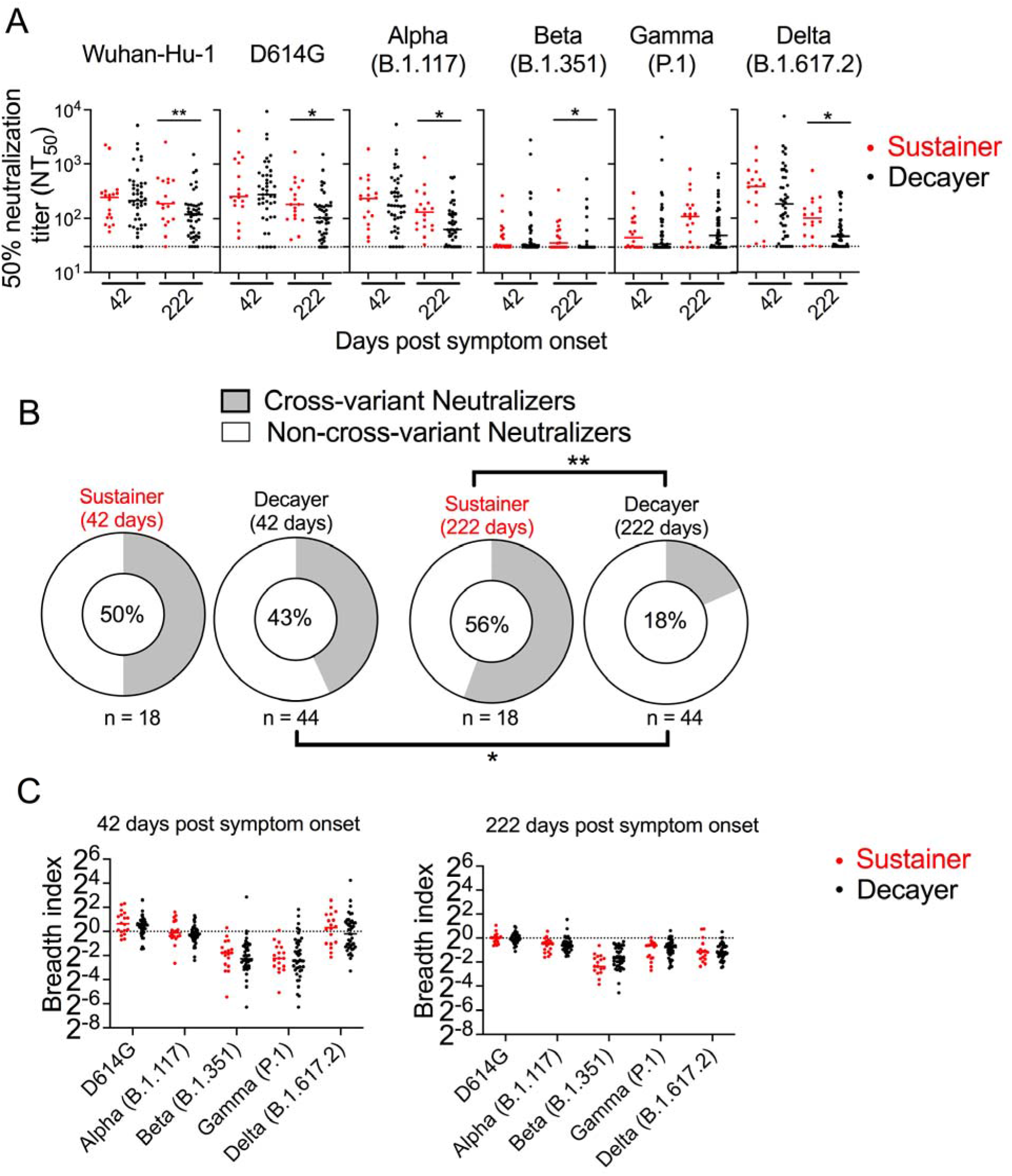
Anti-S IgG sustainers produced superior cross-variant neutralizing capability. (A) Dot plots showing pseudovirus neutralization titers to the indicated SARS-CoV-2 variants in plasma collected at the indicated times (median 42, range 13-119; median 222, range 185-271) from sustainers (n = 18, red) and decayers (n = 44, black). Mann-Whitney U test. (B) Donut plots illustrating percentages of cross-variant neutralizers at the indicated times after symptom onset. Subjects were scored as cross-variant neutralizers if their plasma neutralization titer was more than the detection limit (i.e. 30) for all the variants tested in (A). Fisher exact test. (C) Dot plots showing breadth indexes of plasma collected from anti-S IgG sustainer (red) and decayer (black) at the indicated time after symptom onset. *p< 0.05, **p< 0.01.

When tested early after COVID-19 recovery (~40 days post symptom onset), approximately 50% of the sustainer and decayer plasma samples exhibited cross-neutralization function to all the tested strains (Figure 4A). In contrast, 18% of the decayers and 56% of the sustainers had cross-variant-neutralizing function to all tested strains ~220 days after symptoms onset (Figure 4B). More detailed neutralization analysis suggested decayers had more severe loss of neutralization capacities against the Alpha, Beta, and Delta variants ~220 days after symptom onset (Figure S4 E). Correlation analysis suggested that cross-variant neutralization superiority in sustainers is linked to anti-S and anti-RBD antibody levels (Figure S4, F, G).

There were no differences in neutralization breadth indexes between sustainers and decayers at ~40 or ~220 days post symptom onset (Figure 4C), suggesting that the greater absolute neutralization breadth observed in sustainers at later time points is more likely due to greater antibody magnitudes rather than differences in repertoire composition. These results highlight an important role for anti-S antibody dynamic stability in the durable resistance to emerging SARS-CoV-2 variants.

## DISCUSSION

The immune system harbors the ability to generate long lasting and broadly effective antibody responses against pathogens. Vaccines and vaccination strategies in the past have had variable success in accessing the full extent of antibody durability and breadth available from the immune system. While mRNA vaccination has been an initial strong success, data here demonstrate features of dynamic stability and breadth conferred by an immune response to natural infection that can serve as benchmarks of anti-SARS-CoV-2 immune capability as vaccine strategies continue to mature. How to elicit greater dynamic stability more optimally over time with greater functional breadth across viral variants through vaccination alone are key goals not only to guide ongoing development of strategies for SARS-CoV-2, but in vaccine strategies more broadly.

We charted differential antibody durability and breadth dynamics induced by infection, mRNA vaccination, and a hybrid process of vaccination following infection of an early (i.e., pre-VOC) strain of SARS-CoV-2. Natural infection-induced antibodies, while modest in magnitude, exhibited superior stability and cross-variant neutralization breadth than antibodies induced by a two-dose mRNA regimen spaced 3-4 weeks apart in infection-naïve vaccinees. Pre-existing immunity from natural infection primed mRNA vaccine-induced antibody responses for greater antibody magnitude and dynamic stability than that induced by the two-dose mRNA regimen. In line with our findings, a long-term clinical observation study demonstrated significantly higher risk of Delta-related breakthrough infection, symptomatic diseases and hospitalization in naïve vaccinees compared to naturally infected subjects—and infection plus vaccination afforded additional protection (Sivan Gazit, 2021).

Our observations that improved post-infection antibody durability arises from those with HCoV anti-S2 IgG crossreactivity in both the memory B cell compartment as well as relatively greater early anti-S2 secreted antibody responses, provide insights into accessibility of optimal induction of LLPC development. This is consistent with a model in which recalled high-affinity memory B cells may be more likely to become LLPCs. LLPCs are the sources of long-lived secreted antibodies responsible for months/years-long stable antibody plateau levels and tend to have relatively high levels of SHM (Cortes and Georgopoulos, 2004; Phan et al., 2006; Turner et al., 2021a). The magnitude of the stable virus-specific IgG plateau phase after infection or vaccination is likely proportional to the number of sourcing LLPCs induced at the time of immune challenge. In this light, further understanding the ground rules underlying LLPC fate commitment will be important to guide vaccine strategies to maximize long-term antibody durability.

Sources of LLPCs are GC B cells recruited from either the naïve B cell pool or recall of memory B cells (Akkaya et al., 2020; Turner et al., 2021b; Weisel and Shlomchik, 2017). In the case of recalled immunity, memory B cells with higher affinity toward immunizing antigen are preferred candidates to differentiate into antibody secreting cells in response to a subsequent boosting dose (Mesin et al., 2020). In this context, a greater frequency of more mature, high affinity memory B cells may contribute to increased probabilities of LLPC production. Immune responses early after SARS-CoV-2 infection have been characterized by a memory B cell pool with remarkably low SHM one month after symptom onset but shown to slowly increase months thereafter (Chen et al., 2020; Gaebler et al., 2021; Wang et al., 2021) along with antibody affinity enhancement (Gaebler et al., 2021). It follows that since the SHM/affinity state of memory B cells may influence the probability of LLPC development, the state of memory B cell maturation at the time of recall may be a determinant of antibody durability. In our study, all COVID-19 convalescents who received a vaccination did so ~220 days after onset of their original COVID-19 infection, indicating memory B cell maturity at the time of vaccination more advanced than the memory B cell maturation occurring between 1^st^ and 2^nd^ dose of mRNA vaccine (Alice Cho, 2021). Time will tell whether additional mRNA vaccine boosts months after the second dose will generate more durable anti-SARS-CoV-2 antibody in naïve vaccinees.

It is also possible that immune memory conferred by infection as opposed to vaccination (i.e., intact virus, antigen delivery, antigen persistence, etc.) contributes more effective priming of longer lasting antibody responses. In this regard, we observed here that a group of individuals (sustainers) who had evidence of greater infection-proximal SHM and cross-reactivity in their memory B cell pool exhibited more stable antibody dynamics over time. While this outcome may reflect recruitment of B cells that had matured from prior seasonal HCoV infection in sustainers, the fact that sustainer anti-RBD antibody durability was also apparent suggests that LLPC differentiation preference likely spread beyond recalled memory B cells.

Another determinant of antibody longevity is context of immune challenge. Responses induced by live vaccines (measles, mumps, and rubella) persist for centuries, whereas protein-based vaccines are shorter lasting (Amanna and Slifka, 2010). However, antibody durability is not necessarily linked to the use of live virus as long-lived antibody responses are induced to the human papilloma virus (HPV) vaccine, a non-live viral like particle-based vaccine (Kjaer et al., 2020). This suggests that distinct immunological cues can be engineered in the context of a non-live vaccine to result in the generation of long-lived immune responses.

Our antibody dynamic analysis in naturally infected subjects, naïve vaccinees and COVID-19 vaccinees mirrors the clinical outcome in the real-world breakthrough infection and reinfection analysis (Sivan Gazit, 2021), reinforcing the concept that long-term virus-specific antibody magnitude is a strong immune correlate to SARS-CoV-2 protection. Achieving and maintaining robust magnitudes of antibody production can also have a positive effect on recognition breadth across a diversity of SARS-CoV-2 variants. Antibody magnitude correlates with variant recognition breadth likely because human pre-VOC infection-induced S-specific memory B cell repertoires include a low but reliable frequency of specificities that neutralize broadly across variants of concern (Tong et al., 2021). The frequency of variant S-reactive antibodies within a polyclonal composition may also differ between groups as our measurement of an antibody breadth index normalizes for antibody magnitude. This suggests that vaccines and/or vaccine strategies can be improved to enrich antibody responses with anti-variant neutralization breadth.

Although both infection and vaccination cohorts shared similar pre-VOC SARS-CoV-2 S glycoproteins as antibody targets, polyclonal anti-SARS-CoV-2 variant neutralizing antibody recognition breadth indexes evolved differently over time after natural infection compared to mRNA vaccination. It is possible that replicating intact viruses in natural infection may elicit antibody recognition coverage broader than the coverage elicited by the stabilized (i.e., 2P) version of S in current vaccines. Decreased recognition of the Delta variant after a 2^nd^ vaccine dose suggested a dominant focusing response toward the Wuhan variant at the expense of VOC-recognizing clones. Boosting with strategically designed immunogens in parallel or series may access broader coverage of immune variants (Petrova and Russell, 2018), but may not be as simple as inclusion of diverse VOC Ss (Liu et al., 2021), especially given the likelihood of additional variants yet to emerge. In this regard, understanding how natural infection confers greater anticipatory breadth is an important goal to guide booster S designs.

## Acknowledgments

We thank the study volunteers. We thank Steven C. Harrison for comments on the manuscript and Daniela Weiskopf for the AIM assay peptide pools. We thank Grigoriy Losyev for cell sorting and Aaron Schmidt lab for RBD proteins. This study was supported by NIH grants T32 AI007306 (to Y. Chen) and AI139538 (to D.R.W.). Work in the laboratories of B.C. and D.R.W were funded by the Massachusetts Consortium on Pathogenesis Readiness. D.R.W. acknowledges support from Fast Grant funding for COVID-19 science, the Ragon Institute, and the Mark and Lisa Schwartz and the Schwartz Family Foundation and acknowledges the interest of Enid Schwartz. The schematic plot in figure 2 was made in Biorender.

## Author contributions

Y.C. and D.R.W. conceived, designed, and analyzed the experiments. Y.C., P.T., N.W., A.S.M., A.Z., S.H., and A.G. carried out experiments. T.X., Y. Cai, and B.C. contributed recombinant protein. S.H. and A.S.M. recruited participants, executed clinical protocols. Y.C. and D.R.W. wrote the manuscript with input from all co-authors.

## MATERIALS AND METHODS

### Human subjects

Mass General Brigham Institutional Review Board approved this study and protocol. We recruited participants in Boston area through online advertisement on institutional clinical study website open to public and flyers posted in local hospitals (Chen et al., 2020). Adult volunteers with a history of COVID-19 were recruited between March 2020 and June 2020. As described before (Chen et al., 2020), the eligibility criteria include symptom diagnosis by health professionals and a SARS-CoV-2 positive nasopharyngeal swab RT-PCR test. After enrollment, clinical team identified one subject was not diagnosed by nasopharyngeal swab RT-PCR test, but by Quest Diagnostics antibody test and symptoms consistent with COVID-19 (Chen et al., 2020). Participants self-rated their severity of COVID-19 symptoms with a scale from 1 (very mild) to 10 (very severe). They self-reported their symptom onset date and recovery date, with symptom duration being the difference between the two dates. Sixty-two subjects donated 5 or 6 repeated blood draws until middle of December 2020 before they start getting vaccinated. Demographic information is provided in Table S1. Twenty-eight COVID-19 convalescents continued donating blood after they received BNT162b2 or mRNA-1273 vaccines. Demographic information for these individuals is provided in Table S2.

The subjects participated in the vaccination study were recruited in Boston Area after vaccine emergency use authorizations as described above. Blood draws from participants before vaccination were collected when possible. Participants self-reported vaccination type, vaccination dates, any history of COVID-19 diagnosis and other diseases including immunocompromised status due to medical condition or medication, lung disease, diabetes, and cardiovascular disease. Vaccinated subjects who reported no history of a COVID-19 diagnosis were designated as naïve vaccinees, and this was confirmed with anti-N IgG screening. One participant screened positive for anti-N IgG but was negative for anti-S IgG and anti-RBD IgG within 7 days of the first dose, so was considered a naïve vaccinee. Blood draws collected before or within one day of the 1^st^ dose were designated as pre-vaccination draws. A blood draw for each subject collected after the 1^st^ vaccine dose and before or within 4 days of the 2^nd^ vaccine dose were designated as post 1^st^ dose blood draws. Detailed demographic information describing the naïve vaccinees in this study are provided in Table S3.

### Plasma and PBMC isolation from blood

Blood from participants were collected in EDTA tubes, which were centrifuged at 1200 rpm for 10 min to collect the plasma in the upper layer. Plasma was further spun for 10 min at 2200 rpm to remove debris, and then divided into aliquots to store at −80°C. The lower layer was diluted 1:1 in PBS and loaded on Ficoll (GE lifescience) for PBMCs separation at 2200 rpm for 20 min. PMBCs were washed once in PBS and then resuspended in FBS containing 10% DMSO and aliquoted for −80°C and liquid nitrogen storage.

### Cell lines

HEK293T cell line (ATCC) was maintained in DMEM containing 10% FBS, 100 U/mL penicillin and 100 U/mL streptomycin (Gibco). HEK293T stable cell line overexpressing human ACE2 and TMRPSS2 (ACE2/TMPRSS2-expressing HEK293T) was kindly provided by Dr. Marc Johnson (University of Missouri School of Medicine). All cell lines were incubated at 37 °C with 5% CO_2_.

### Recombinant coronavirus proteins

SARS-CoV-2 RBD was kindly provided by Aaron Schmidt’s lab (Ragon Institute of MGH, MIT and Harvard). SARS-CoV-2 Nucleocapsid was purchased from Genscript (Z03480). SARS-CoV-1 spike (40634-V08B), MERS spike (40069-V08B), HKU1 spike (40606-V08B), OC43 spike (40607-V08B), NL63 spike (40604-V08B), 229E spike (40605-V08B), SARS-CoV-2 NTD (40591-V49H) and S2 (40590-V08B) proteins were purchased from Sino Biological.

### Generation of plasmids expressing SARS-CoV-2 variants

We designed 5’ and 3’ primers specific for each variant (Alpha, Beta, Gamma, Delta, each supplied by the Bing Chen laboratory), and amplified the variant sequences by Phusion^®^ High-Fidelity DNA Polymerase (Thermo fisher). We cloned each variant into pHDM vector through EcoRI and SalI restriction enzyme cutting site. To be consistent with the pHDM-Wuhan-Hu-1-delta21 and pHDM-D614G-delta21 plasmids obtained from Addgene, the c-terminal 21 amino acid on all the variants were deleted (Crawford et al., 2020).

### Flow cytometry and single cell sort

Frozen PBMCs were quickly thawed at 37°C and transferred to warm RPMI containing 10% FBS. S^+^ memory B cells were single sorted (BD FACSAria™ Fusion) into 96 well PCR plates as described before (Chen et al., 2020). Briefly, B cells were enriched by anti-CD19 magnetic beads (Miltenyi) and flow through was collected for T cell analysis. Enriched B cells were stained with Flag tagged SARS-CoV-2 spike (Genscript, Z03481) then incubated with APC conjugated anti-Flag and PE conjugated anti-Flag for double staining. DAPI^-^IgM^-^IgD^-^ IgG^+^CD27^+^Spike^+^ cells were single sorted into 96 well PCR plates with 4μL/well lysis buffer (0.5x PBS containing 10mM DTT, and 4 U RNaseout) and stored at −80°C for further analysis (Tiller et al., 2008).

### Antibody sequencing, cloning and expression

RNA from single-sorted S^+^ memory B cell were reverse transcribed into cDNA followed by IgH, Igκ, and Igλ gene amplification through semi-nested PCR as described (Chen et al., 2020; Tiller et al., 2008). The PCR products from second amplification were sent for Sanger sequencing. Sequences were filtered through GENEWIZ-provided quality score, then aligned by IgBlastn v.1.16.0. V gene segment-specific forward primers and J gene segment-specific reverse primers were used in the PCR reactions to amplify specific antibody genes from the 1^st^ round PCR products. PCR products were cut with restriction enzymes, purified, then ligated into expression vectors (Tiller et al., 2008). Ligation products were then transformed in *E.coli* Top10 competent cells (Thermo fisher) to generate colonies which were picked for plasmid extraction and sanger sequencing. Once plasmid sequences were confirmed through Igblast and aligned with the original 2^nd^ round PCR product sequences, paired IgH and IgL plasmids were transfected into 293T cells by lipofetamine 3000 (Thermo fisher). Supernatants from 293T cells were harvested 6 -7 days after transfection and screened in the ELISA for IgG expression. Supernatants with IgG expression were purified by protein A (Thermo fisher), eluted in 150 μL 0.1 M glycine (pH 2.8), and collected in 15 μL 1 M Tris (pH 8.0) (Ho et al., 2016; Tiller et al., 2008). Elution was further dialyzed in sterile PBS every three hours for three times based on manufacturer’s instructions (Thermo fisher, 88260). Concentrations of purified antibody were measured by Nanodrop (Thermo fisher), and stored at 4 °C.

### ELISA

Plasma IgG quantification of coronavirus antigens were performed as previously described (Chen et al., 2020). Briefly, 96 well maxisorp ELISA plates (Thermo fisher) were coated with 2 μg N protein, 2 μg S protein, or 2 μg RBD protein in 30μL PBS overnight at 4°C. After discarding coated solutions, ELISA plates were blocked in 100 μL 3% BSA for 2 hours. During the incubation time, plasma was transferred to tubes and combined with equal volume of 2% Triton in PBS and incubated 20 min at room temperature for inactivation. Plasma from COVID-19 convalescents were serially diluted 2 times from 1:100 to 1:6400, and post-vaccination plasma samples were serial diluted 3 times from 1:100 to 1:72900 in PBS containing 1% BSA and 0.05% Tween20. For the anti-S and anti-RBD antibody standard, the monoclonal antibody CR3022 was serially diluted 2 times from 0.5 μg/mL. After blocking solution was discarded, ELISA plates were washed once in PBS containing 0.05% Tween20. The plasma dilutions were transferred to ELISA plates with duplicates and standards included on each plate as controls. After overnight incubation at 4 °C, ELISA plates were washed three times in PBS containing 0.05% Tween20, and then incubated 90 min with alkaline phosphatase (AP) conjugated anti-human IgG (Southern biotech) diluted at 1:1000 in PBS containing 1% BSA and 0.05% Tween20. ELISA plates were washed three times in PBS containing 0.05% Tween20, and then incubated with 100 μL alkaline phosphatase substrate solution (Sigma) for 2 hours. Plates were read at 405 nm through a microplate reader (Biotek Synergy H1). Graphpad software generated standard curves for each plate by non-linear regression. The plasma Ig level was interpolated from a single dilution with OD_405_ values falling in the middle range of the standard curve. Antibody binding values were reported as CR3022 monoclonal antibody concentration equivalents (mAb μg/mL equivalent). Four pre-COVID controls were included on each plate. Samples were scored positive if the antibody level was higher than 2x average of the pre-COVID controls for each antigen. Diluted plasma from serial draws of each subject were placed on the same ELISA plate. For longitudinal samples measured on different plates, samples from prior draws were rerun to calibrate the experimental variances across plates. For trajectory analysis, one phase decay model and linear regression model of log transformed antibody data were assessed in all cases (Dan et al., 2021). An extra sum-of-squares F Test was used for fitting curve comparisons. One phase decay model was used to report R, peak and plateau anti-S and anti-RBD IgG using log transformed data. Linear regression was used to report slope.

To test mAb binding to other coronavirus spike, ELISA plates were coated with 2 μg SARS spike, MERS spike, HKU1 spike, OC43 spike, NL63 spike or 229E spike in 30 μL PBS. To test mAb binding to SARS-CoV-2 subdomains, ELISA plates were coated with 4 μg SARS-CoV-2 S1, 2 μg SARS-CoV-2 NTD, 2 μg SARS-CoV-2 RBD, and 2 μg SARS-CoV-2 S2. Coated plates were incubated at 4 °C overnight, and then blocked in 3% BSA as described above. Plates were then incubated with 1 μg/mL mAb in PBS containing 0.05% Tween20 at 4 °C overnight. ELISA plates were washed and incubated with secondary antibody as described above. After plates were washed three times, ELISA plates coated with other coronavirus spike were incubated with 100 μL alkaline phosphatase substrate solution overnight, and ELISA plates coated with SARS-CoV-2 subdomains were incubated with substrate solution 2 hours. All plates were read at 405 nm through a microplate reader (Biotek Synergy H1). Binding signals were scored positive if OD_405_ values were above 5x PBS control on the same plate.

### mAb spike binding EC_50_ measurement

293T cells were co-transfected with pHDM-Wuhan-Hu-1 and pMAX-GFP plasmids at a 4:1 ratio by lipofectamine 3000 (Thermo fisher). 293T cell pool cotransfected with pHEF-VsVg and pMAX-GFP plasmids at a 4:1 ratio was used as a negative control. Cells were harvested 2 days post transfection and incubated with serially diluted mAbs. A total of nine three-fold serial dilutions were performed for each mAb from 10 μg/mL. These dilutions were incubated with 2×10^5^ transfected 293T cells on ice for 20 min. Cells were washed and stained with DAPI and AF647-anti-human IgG (Thermo fisher, A-21445), and analyzed by flow cytometry (BD Canto II). GFP^+^ cells were gated to measure MFI (AF647 fluorescence) of S-expressing 293T cells. If MFI of 10 μg/mL mAb-treated pHDM-Wuhan-Hu-1/pMAX-GFP-co-transfected 293T cells at was at least 2 times greater than mAb treated negative control cells (pHEF-VsVg/pMAX-GFP cotransfected 293T cells), the mAb was scored as an S binder. The EC_50_ for each S-binding mAb was reported as the concentration with 50% of maximal binding determined by non-linear regression using Graphpad Prism. If the maximal binding of the mAb was not achieved at the concentration 10 μg/mL, EC50 was set as the detection limit (i.e. 10 μg/mL).

### Activation induced markers (AIM) assay

After 2~3×10^6^ PBMCs were enriched for B cells through anti-CD19 magnetic beads (Miltenyi), flow through cells were centrifuged and incubated with 1 μg/mL peptide megapools consisting of SARS-CoV-2 spike, SARS-CoV-2 remainder (SARS-CoV-2 proteome minus spike), HCoVs (HKU1, OC43, NL63, 229E) spike, HCoVs remainder (HCoVs proteome minus spike), and cytomegalovirus (CMV), as well as phytohemagglutinin (500x, Thermo fisher) and DMSO. Cells were cultured in 96 well round-bottom plates with RPMI containing 10% FBS for 24 hours. Cells were stained with Biolegend antibodies FITC anti-CD27 (356404), PE anti-CCR7 (353204), Percp cy5.5 anti-CD4 (357414), PE-cy7 anti-CD8 (344712), APC anti-OX40 (350008), APC-Cy7 anti-CD45RA (304128), AF700 anti-CD3 (300424), BV510 anti-CD19 (302242), BV510 anti-CD14 (367124), BV605 anti-CD69 (310938), BV711 anti-CD137(309832).

### SARS-CoV-2 pseudovirus neutralization assay

293T cells were co-transfected with pHDM vector expressing SARS-CoV-2 variants, pLenti CMV Puro LUC (w168-1), and psPAX2 (Addgene) with lipofetamine 3000. Pseudoviruses in the supernatants were harvested two days post transfection, and diluted in DMEM containing 10% FBS to test titers in 293T cells stably expressing human ACE2 and TMRPSS2 (provided by Dr. Marc Johnson, University of Missouri School of Medicine). For neutralization assays (Crawford et al., 2020), plasma was serially diluted to incubate with pseudovirus at 37°C for 1 hour. The mixtures of diluted plasma and pseudovirus in duplicate were added to 293T cells stably expressing human ACE2 and TMRPSS2 (2×10^4^/well). Two days after the incubation, cells were lysed by Promega One-Glo luciferase reagent, and the luciferase signals were measured by Biotek Synergy H1. Plasma titers that achieved 50% neutralization (NT_50_) were determined by non-linear regression using Graphpad prism of log-transformed luciferase signal. For early neutralization response analysis, the second-month blood draw (instead of the first draw) of 9 COVID-19 convalescent subjects was used in neutralization test to Wuhan-Hu-1, D614G, Alpha, Beta, Gamma and Delta variants because of depletion of the first month plasma from these 9 subjects. Unless the lowest plasma dilution (1:30) returns NT_50_ lower than 50%, NT_50_ derived from non-linear regression curve with goodness of fit less than 0.7 was excluded. NT_50_ was set as the limited of detection (i.e. 30) if the first plasma dilution (1:30) returned less than 50% neutralization (Chen et al., 2020). Plasma samples that showed Wuhan-Hu-1 strain NT_50_ and variant NT_50_ values no more than 30 (detection limit) were excluded from breadth index calculations.

### Statistical analysis

Statistical analysis was performed with Graphpad prism 8 software. After normality testing, log-transformed antibody titers were compared by student’s *t* test for two comparisons and ANOVA with Tukey’s test for multiple comparisons. Unless indicated, two tailed Mann-Whitney U tests were performed to analyze statistical differences between two comparisons; Two tailed Kruskal-Wallis test with Dunn’s multiple comparison correction was performed to analyze statistical differences in multiple comparisons. Spearman correlation was used for association analysis.

Extra sum-of-squares F Test was used for comparisons of fitting curves. Unless indicated, horizontal lines in dot plots indicate median. Analysis of Covariance (ANCOVA) was performed to test for significance of differences in slopes generated from simple linear regression.

## KEY RESOURCES TABLE

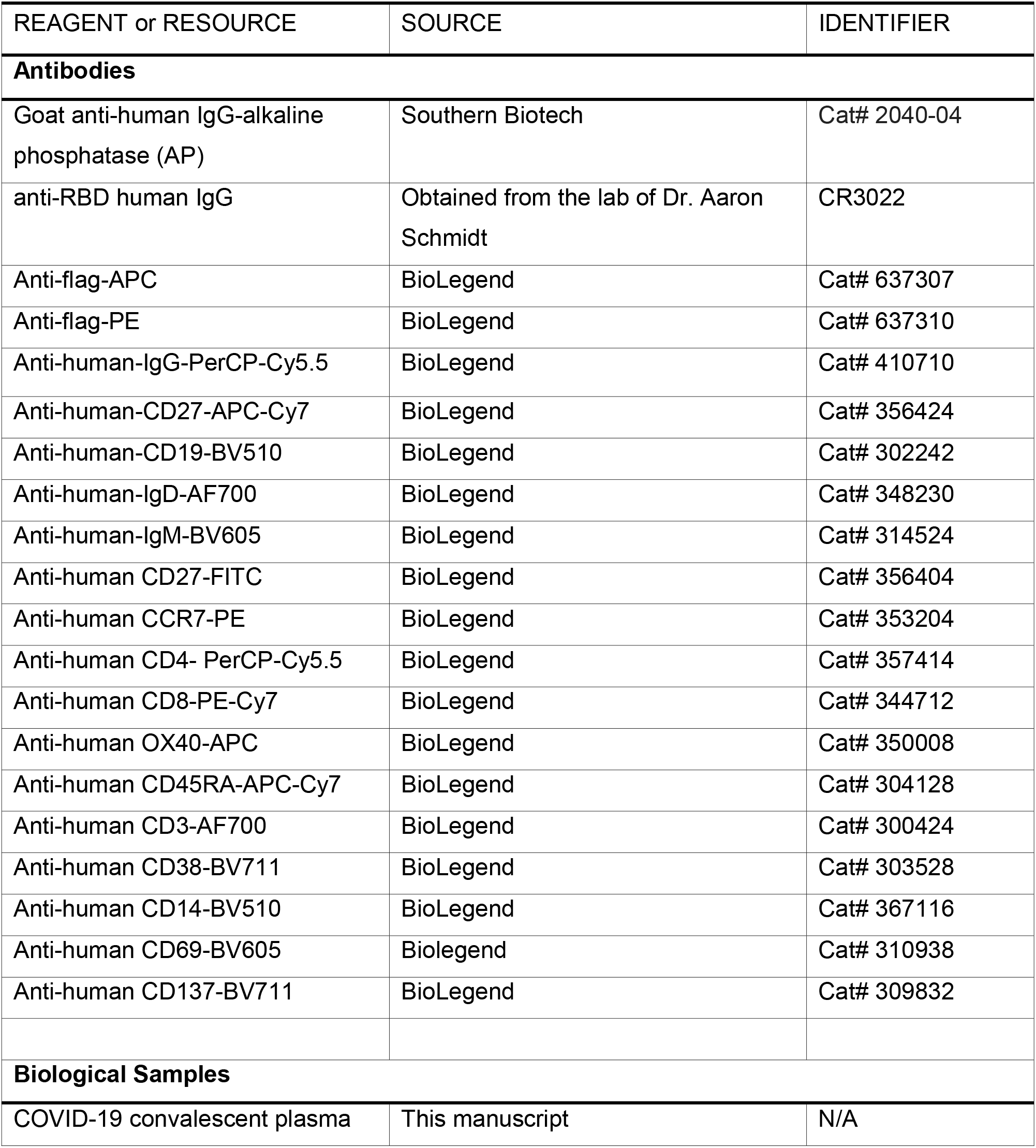
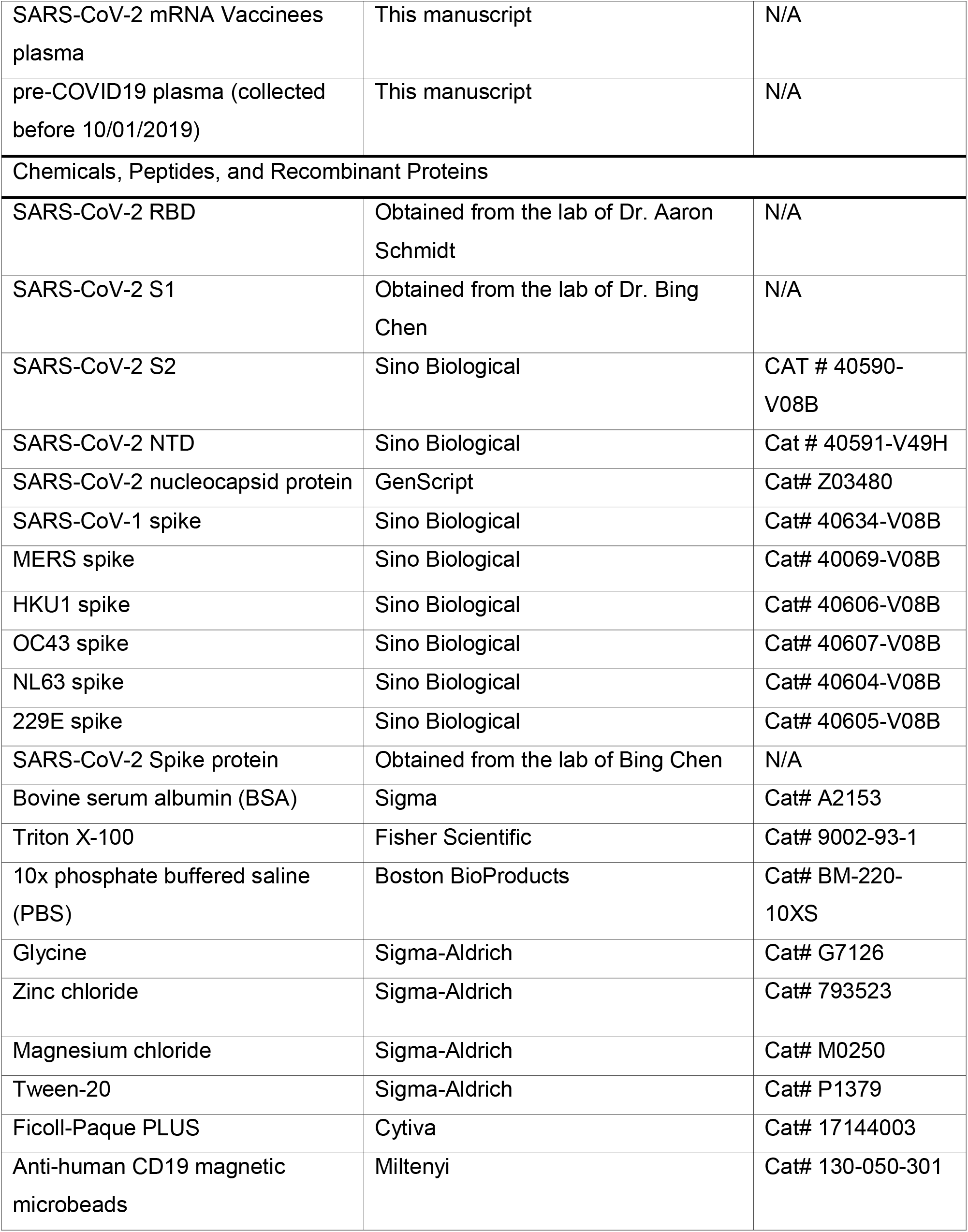
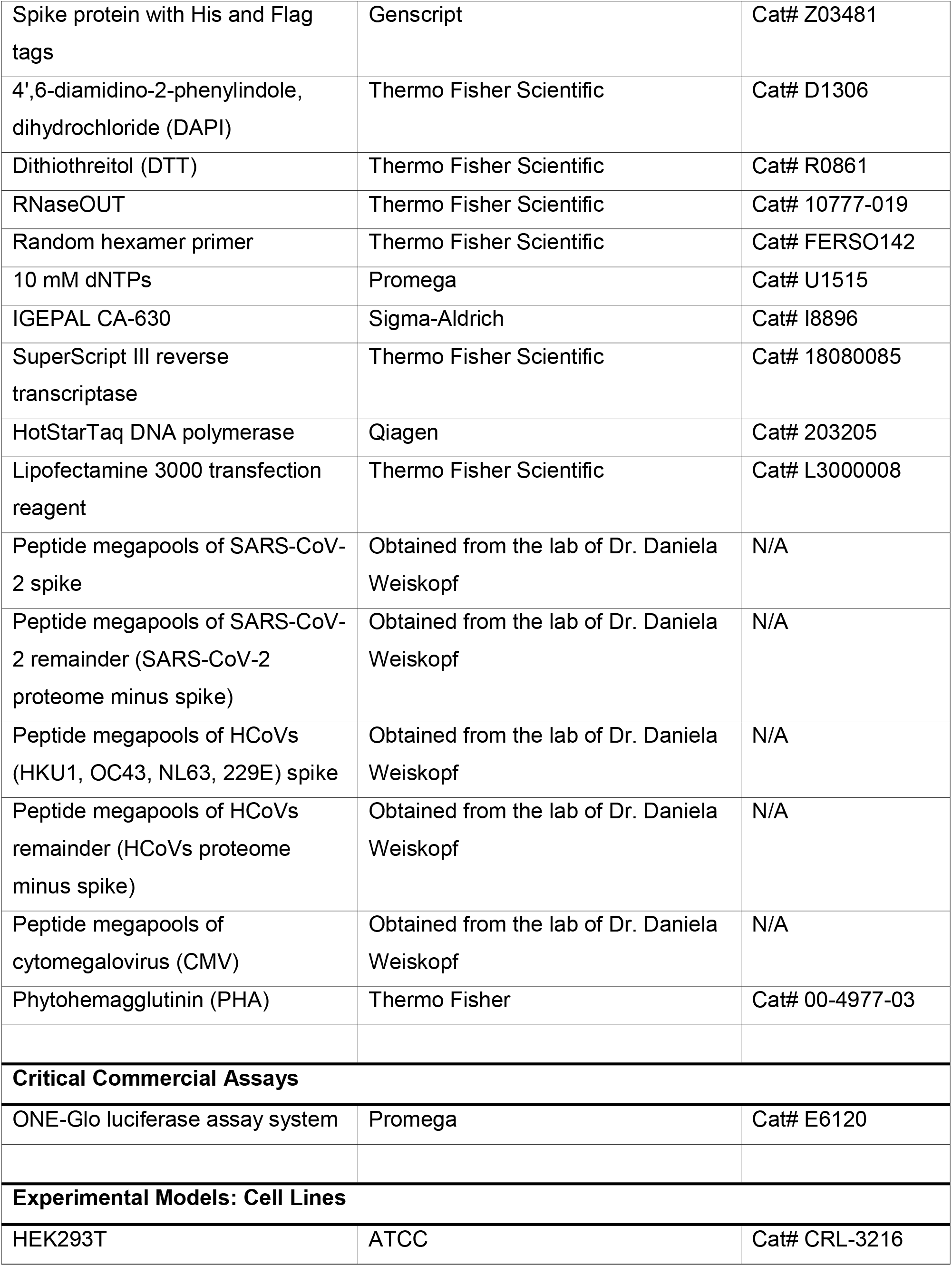
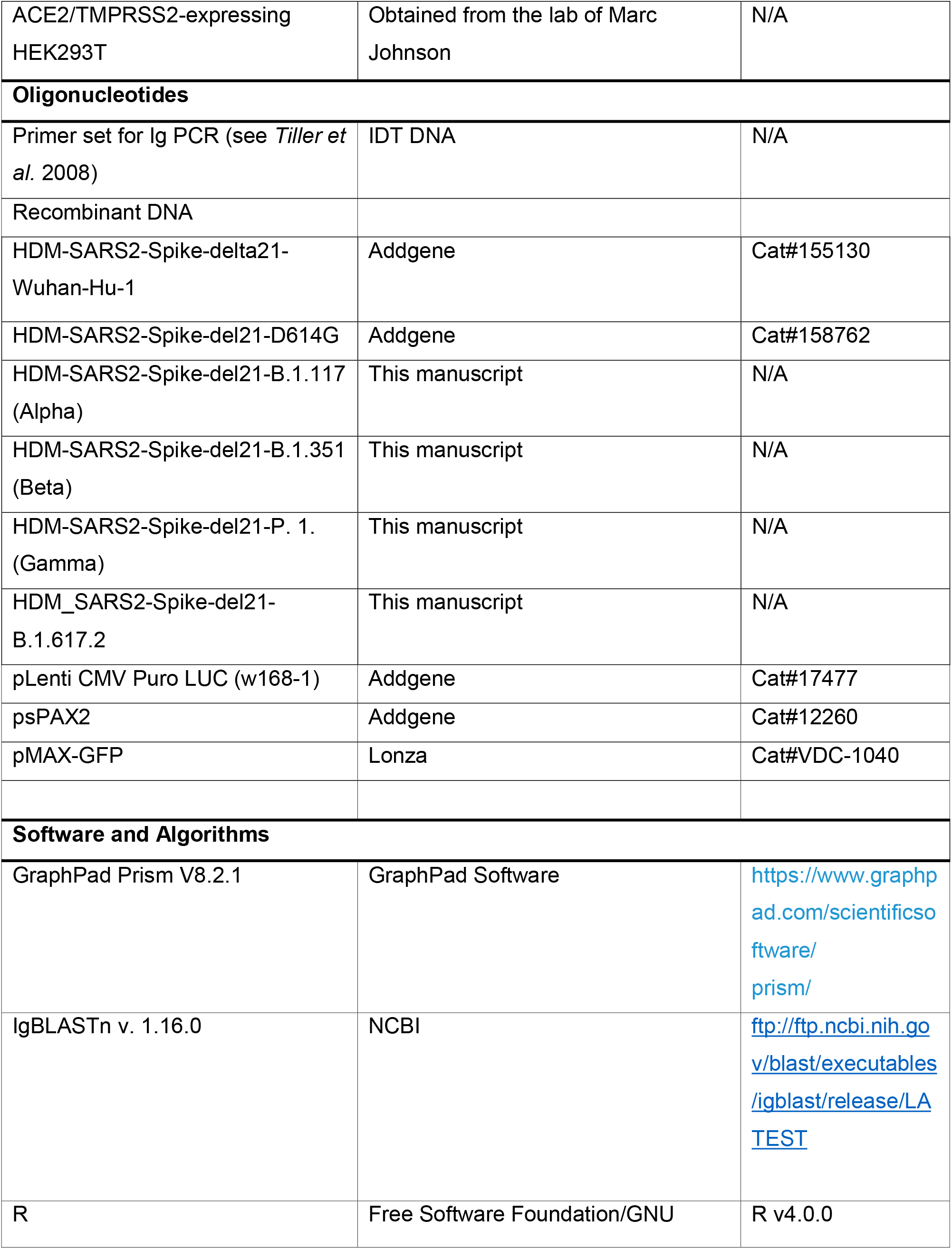
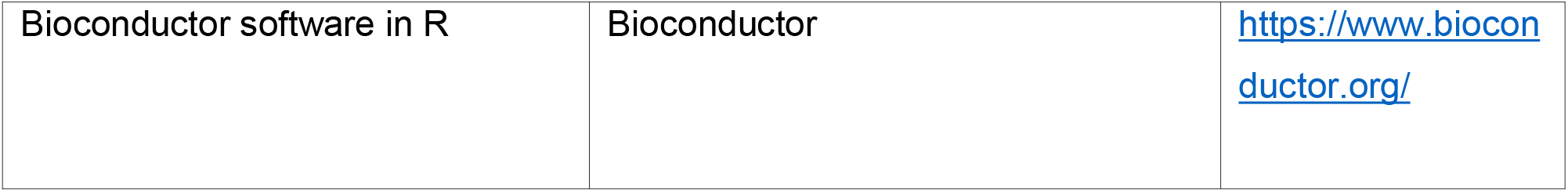

---

**Figure S1.**
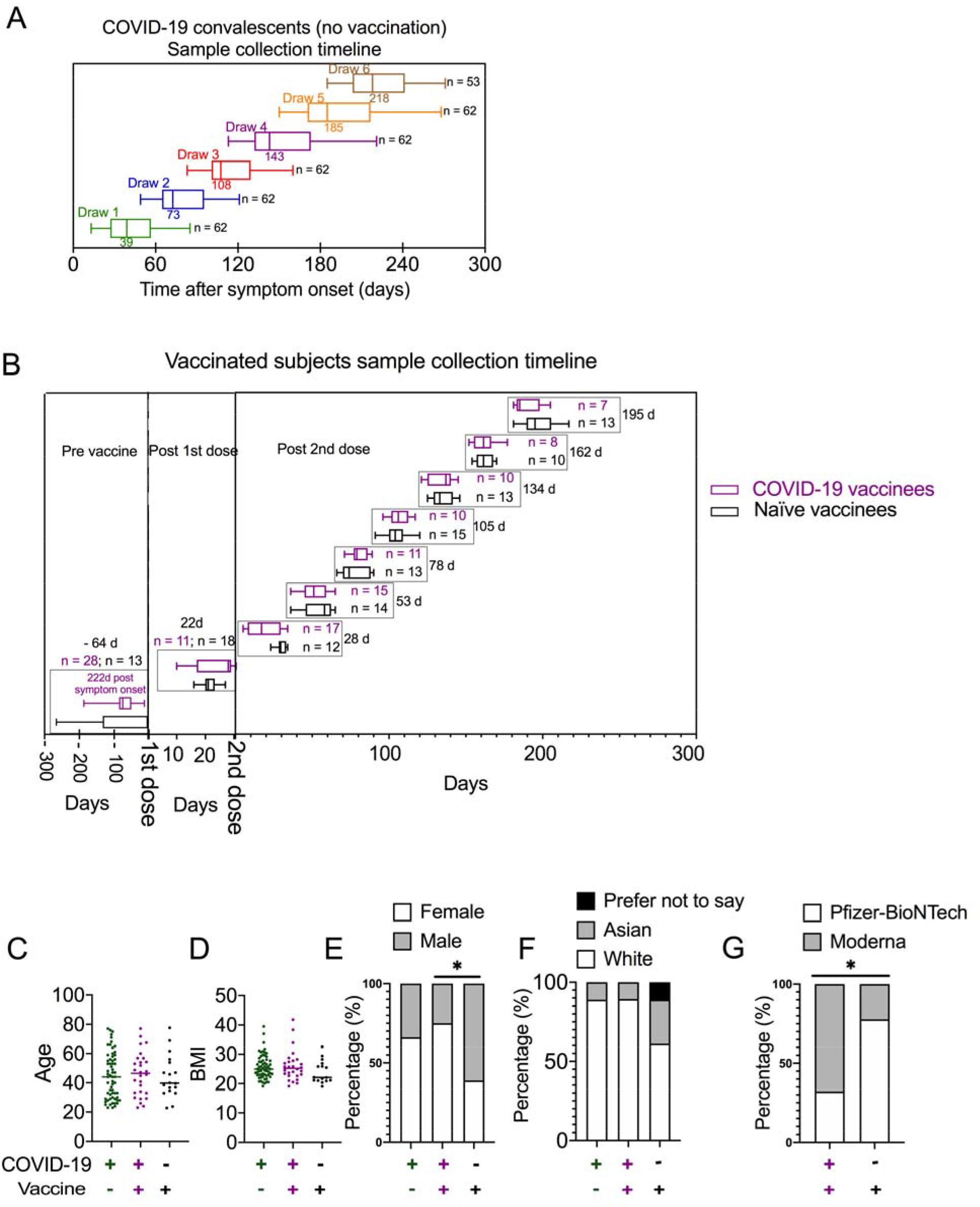
Sample collection timeline and cohort characteristics. (A) Box and whisker plots showing blood collection timeline for seroconverted COVID-19 convalescent subjects before vaccination. 62 and 53 subjects donated 5 and 6 repeated blood draws, respectively, before vaccination. Median time is indicated. (B) Box and whisker plots showing blood collection timeline for COVID-19 vaccinees (purple, n = 28) and naïve subjects (black, n = 18) with mRNA vaccination. The x-axis shows days before 1^st^ dose, after 1^st^ dose, and after 2^nd^ dose. Median days and number of subjects donating blood in each time interval are indicated. (C-G) Dot and bar graphs of age (survey 100% complete), BMI (survey 94% complete), gender (survey 100% complete) and race (survey 89% complete) distributions in COVID-19 convalescent subjects before vaccination, COVID-19 vaccinees, and naive vaccines as described in (A, B). (G) Bar graph showing vaccine type distribution in COVID-19 vaccinees, and naive vaccines as described in (B). Kruskal-Wallis test was performed for statistical analysis in C and D. Fisher exact test was performed for E-G. *p< 0.05.

**Figure S2.**
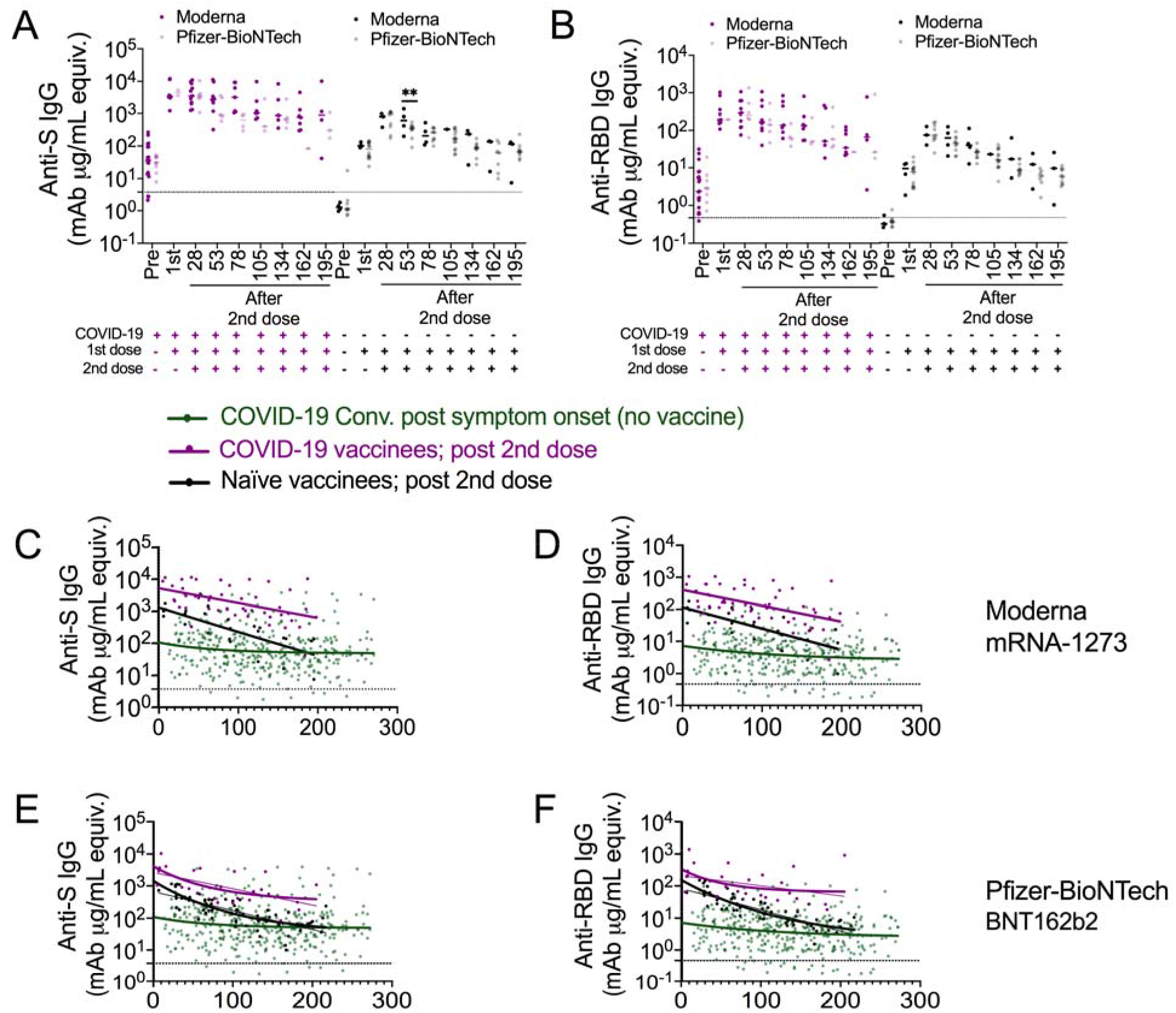
Antibody response comparisons between Moderna mRNA-1273 and Pfizer-BioNTech BNT162b2 vaccines. (A, B) Dot plots showing plasma anti-S (A) and anti-RBD (B) IgG level in COVID-19 vaccinees (purple, n = 4-19) and naïve vaccinees (black, n = 1-4) receiving Moderna mRNA-1273 vaccination, and COVID-19 vaccinees (light purple, n = 1-9) and naïve vaccinees (grey, n = 7-14) receiving Pfizer-BioNTech BNT162b2 vaccination at the indicated median collection time point (days). (C-F) Dot and line graph showing anti-S (C, E) and anti-RBD (D, F) IgG trajectory over time with one phase decay model (thick line) and linear regression (thin line) fitted curves in COVID-19 convalescents (green), as well as COVID-19 vaccinees (purple) and naïve vaccinees (black) receiving Moderna mRNA - 1273 vaccination (C, D), and Pfizer-BioNTech BNT162b2 mRNA vaccination (E, F).

**Figure S3.**
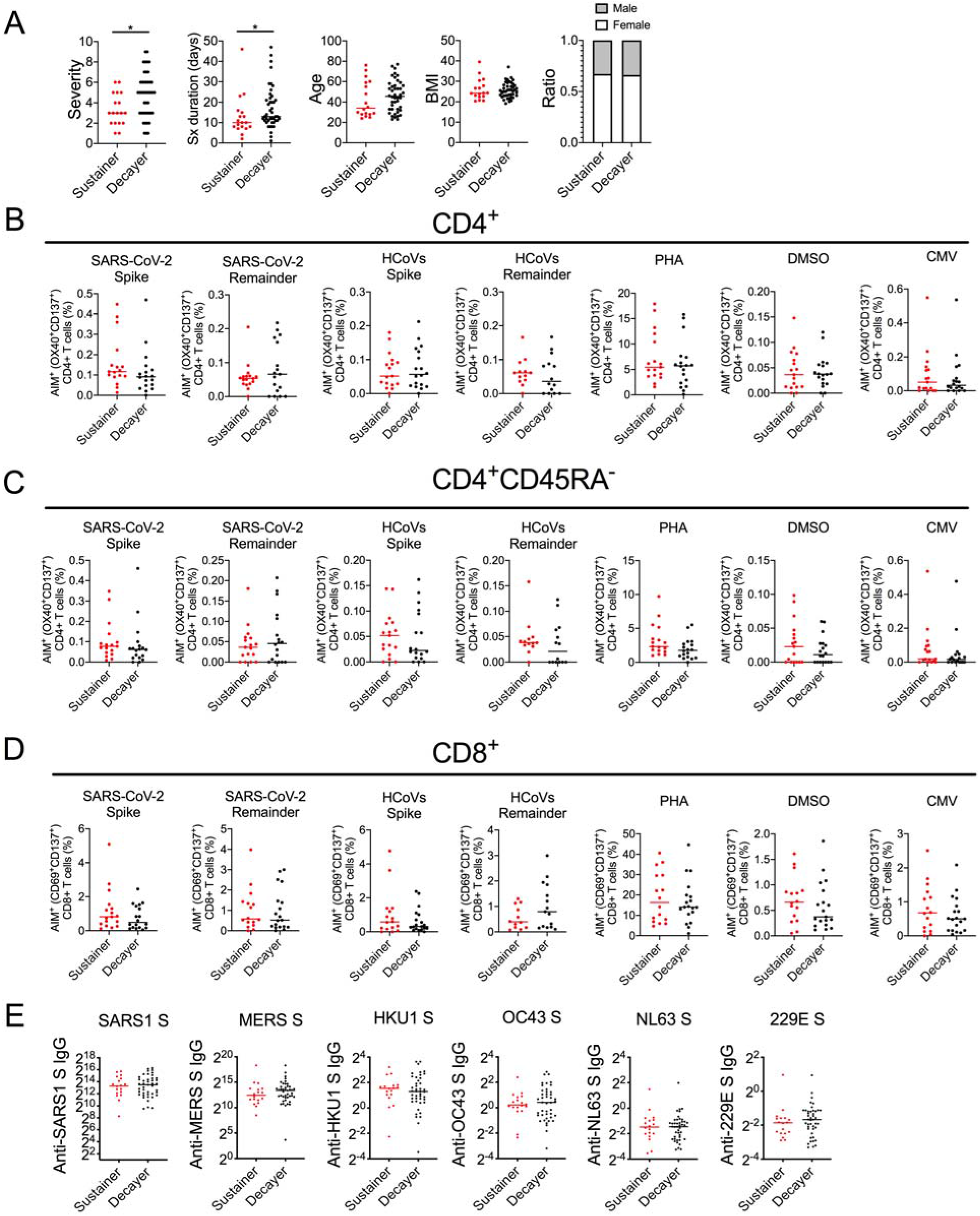
T cell and plasma cross-reactivity analysis in sustainers versus decayers after natural infection. (A) Plots showing the distributions of symptom duration, severity, age, BMI, and genders between sustainers (red, n = 18) and decayers (black, n = 44). Surveys were more than 94% complete for each category. Mann-Whitney U test. (B-D) Dot plots showing the percentage of AIM^+^(OX40^+^CD137^+^) cells (B, C) and AIM^+^ (CD69^+^CD137^+^) cells (D) gated on live CD4^+^ (B), CD4^+^CD45RA^-^ (C), and CD8^+^ (D) cells as indicated in sustainers (n = 13-17) and decayers (n = 14-18) after day ~40 post-symptom-onset PBMCs were stimulated with SARS-CoV-2 and seasonal HCoVs peptide megapools consisting of full length spike or non-spike peptides remaining, termed remainders (i.e., whole proteome minus spike). Mann-Whitney U test. (E) Dot plots showing HCoV spike-specific IgG in plasma collected in sustainers (n = 18) and decayers (n = 43) ~40 days after symptom onset. Mann-Whitney U test. *p<0.05.

**Figure S4.**
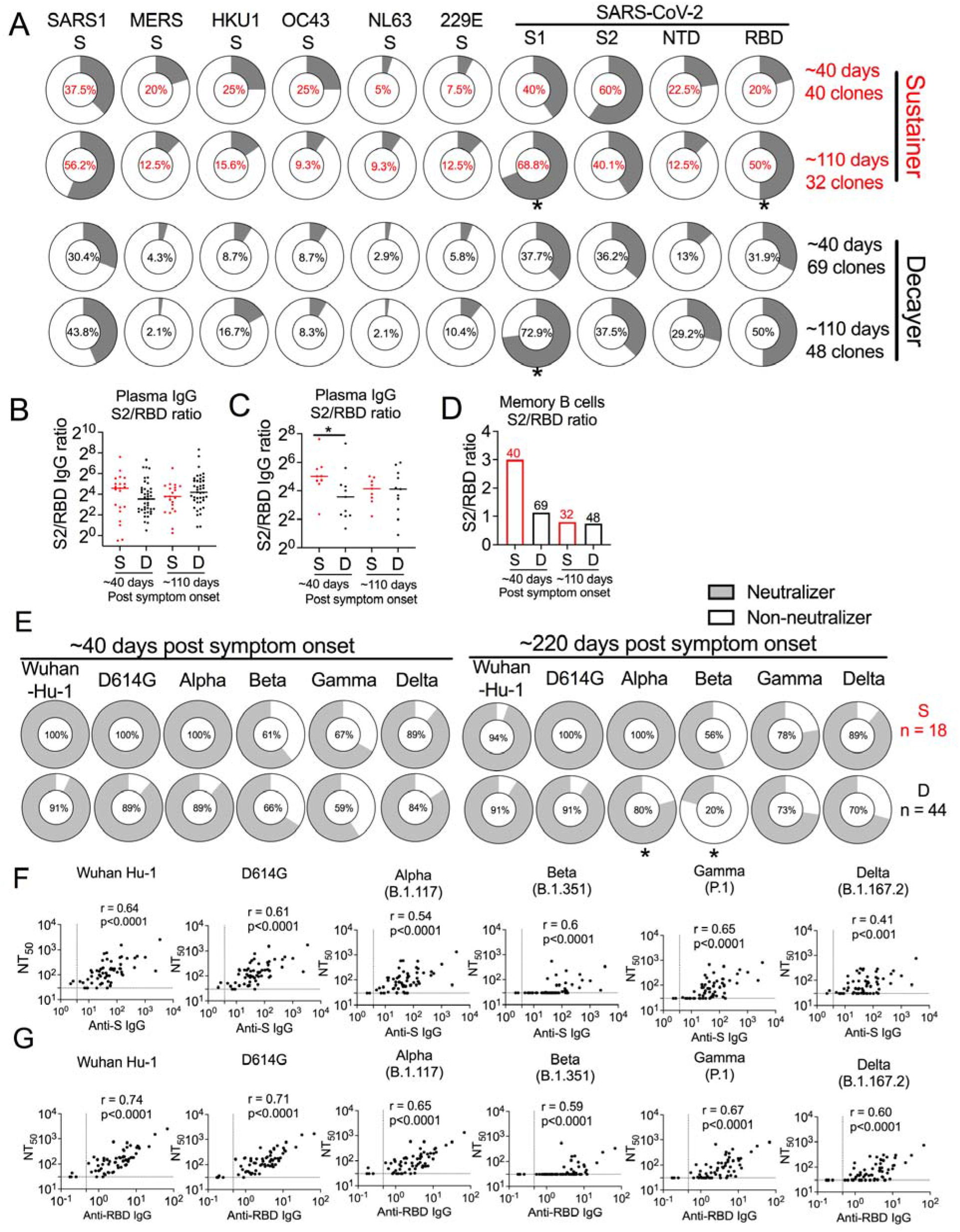
Evolution of antibody recognition of coronaviruses and analysis of neutralization function in sustainers versus decayers after natural infection. (A) Dot plot showing the binding frequency of sustainer-isolated (red) versus decayer-isolated (black) S^+^ memory B cell-derived mAbs to the indicated HCoV spike proteins. Memory B cells were collected ~ 40 days after symptom onset (sustainer, n = 9, 40 clones; decayer, n = 11, 69 clones;1^st^ draw) and ~110 days after symptom onset (sustainer, n = 8, 32 clones; decayer, n = 9, 48 clones; 3^rd^ draw). Fisher exact test. (B) Dot plot showing the ratio between plasma anti-S2 IgG and anti-RBD IgG from all anti-S IgG or anti-RBD IgG sustainers (n = 21) and decayers (n = 41) at ~40 or ~220 days after symptom onset. (C) Dot plot showing the ratio between plasma anti-S2 IgG and anti-RBD IgG from sustainers (n = 9) and decayers (n = 11) selected for memory B cell sorting at ~40 or ~220 days after symptom onset. Selection was based on similar initial antibody levels. Mann-Whitney U test. (D) Bar graph showing the ratio of S2-binding mAb frequencies to RBD-binding mAb frequencies in sorted memory B cells from sustainers (S, red) and decayers (D, black) at ~40 or ~220 days after symptom onset. Number of mAbs in this analysis are indicated on the Bar graph. (E) Donut plots illustrating the percentage of neutralizers harboring neutralization titer > 30 (detection limit) to the indicated pseudotyped variants in sustainers (n = 18) and decayers (n = 44) ~40 days (left) and ~220 days (right) after symptom onset. Fisher exact test. (F-G) Scatter plots illustrating Spearman correlation between anti-S (F) and anti-RBD (G) IgG in plasma isolated ~220 days after symptom onset from COVID-19 convalescents (n = 62) and neutralization titers to the indicated variants. *p<0.05, **p<0.01, ***p<0.005, ****p<0.0001.

**Supplemental table 1:**
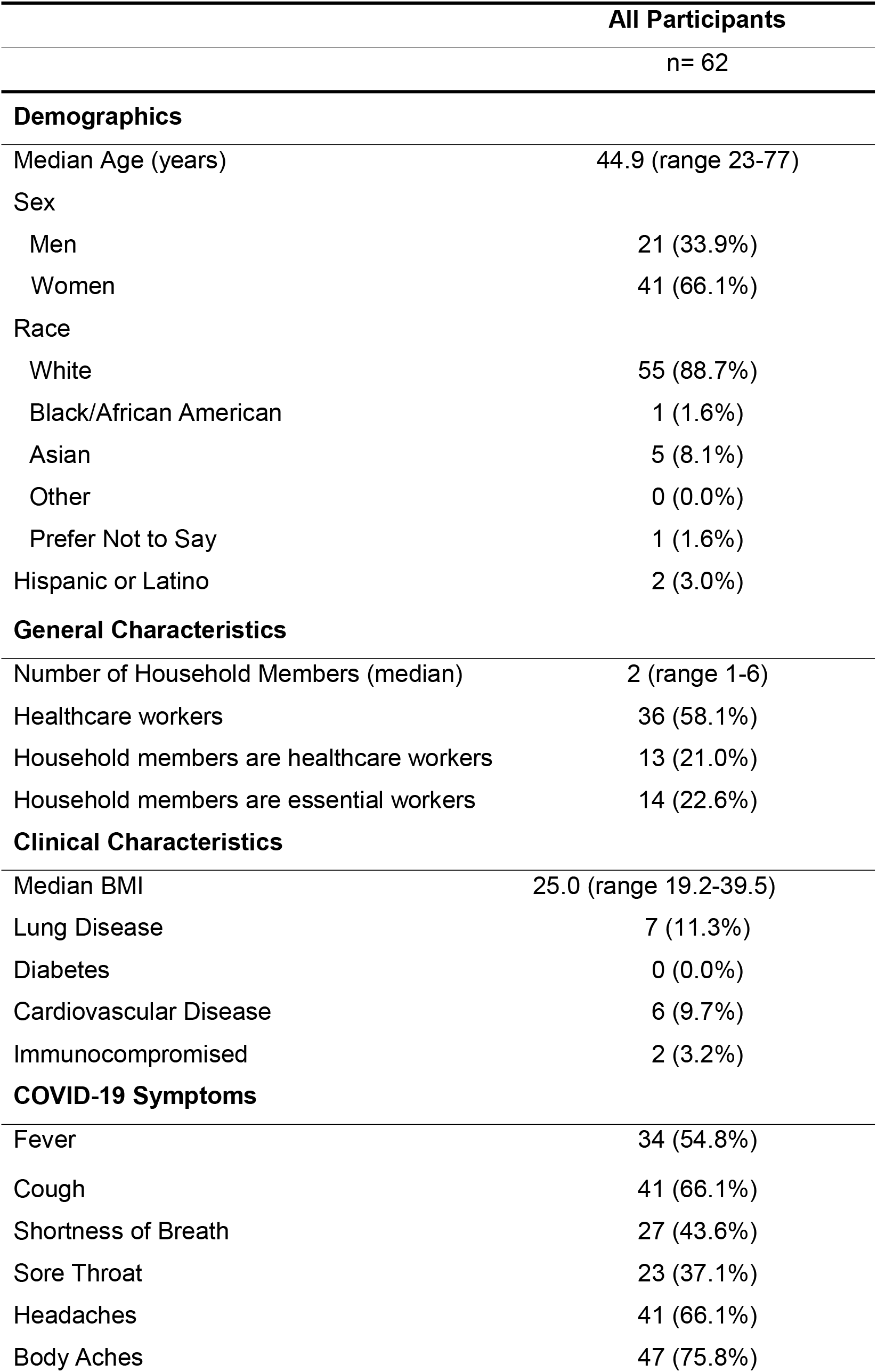
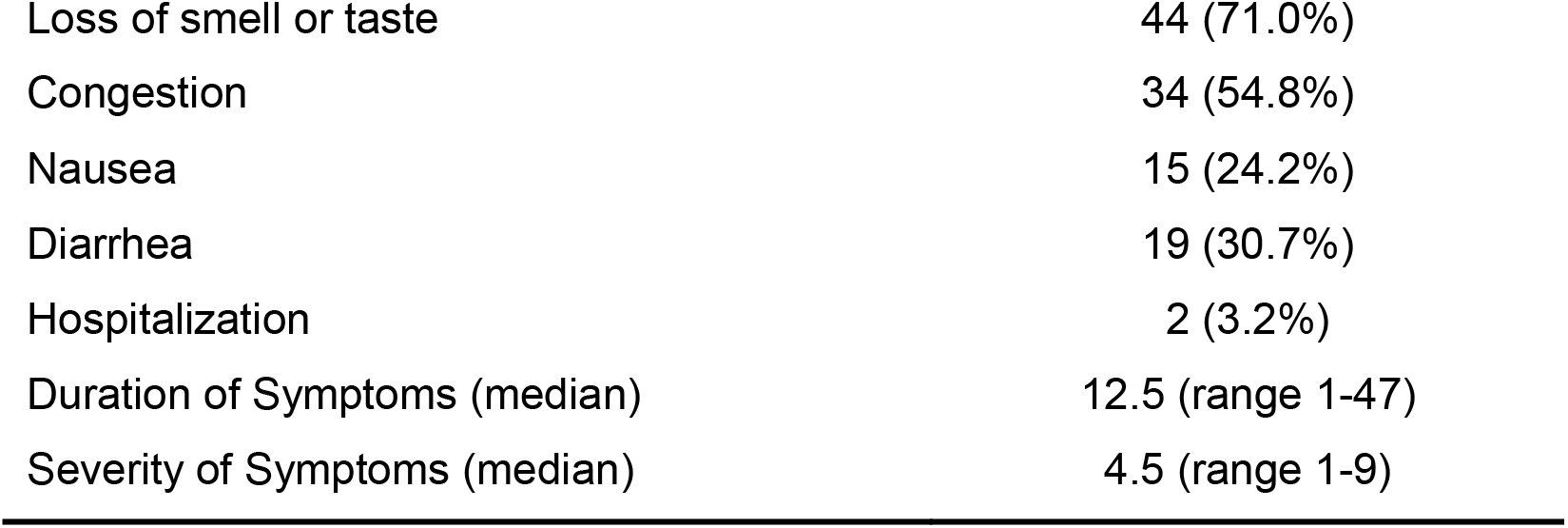
COVID-19 convalescent cohort characteristics.

**Supplemental table 2:**
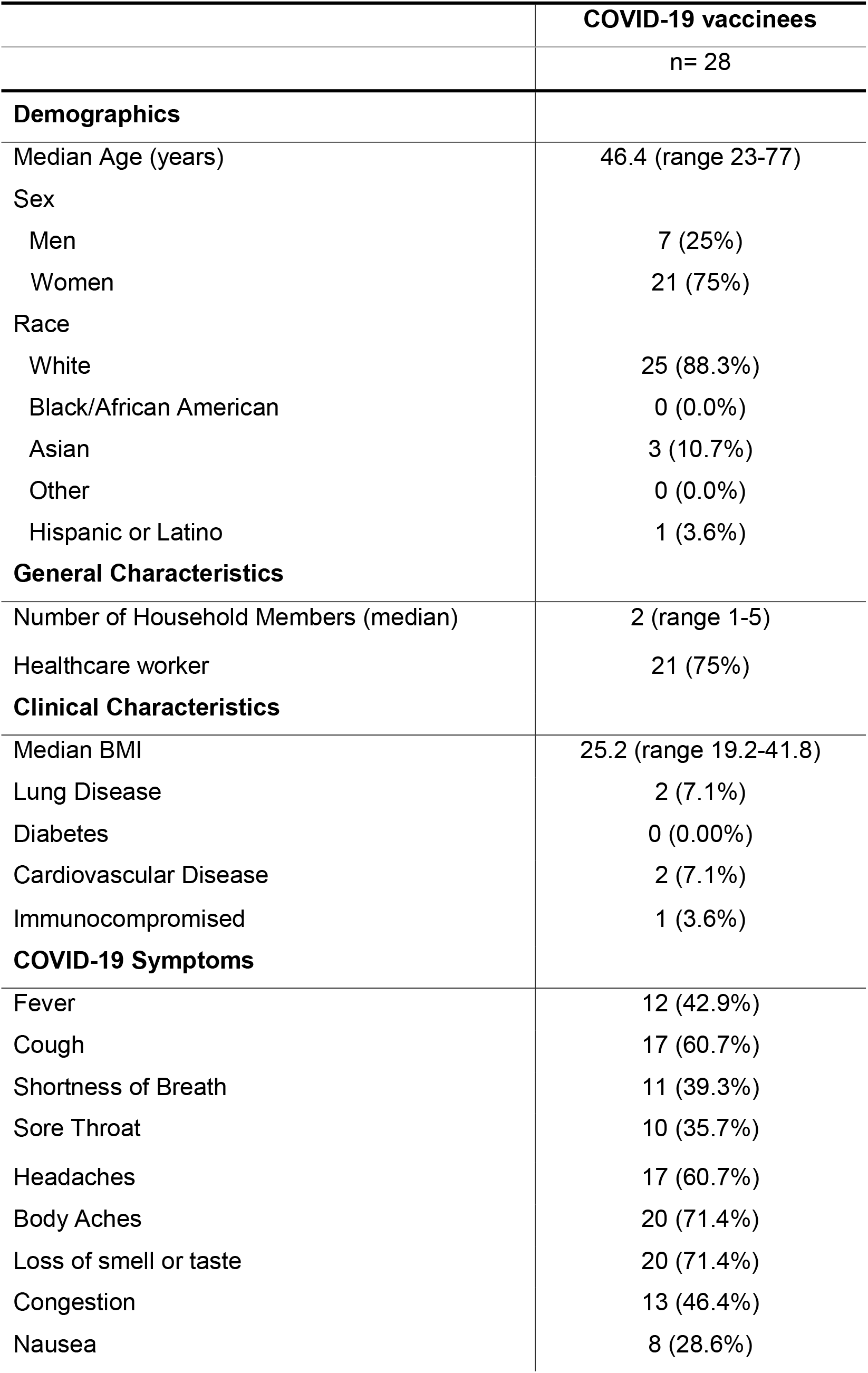
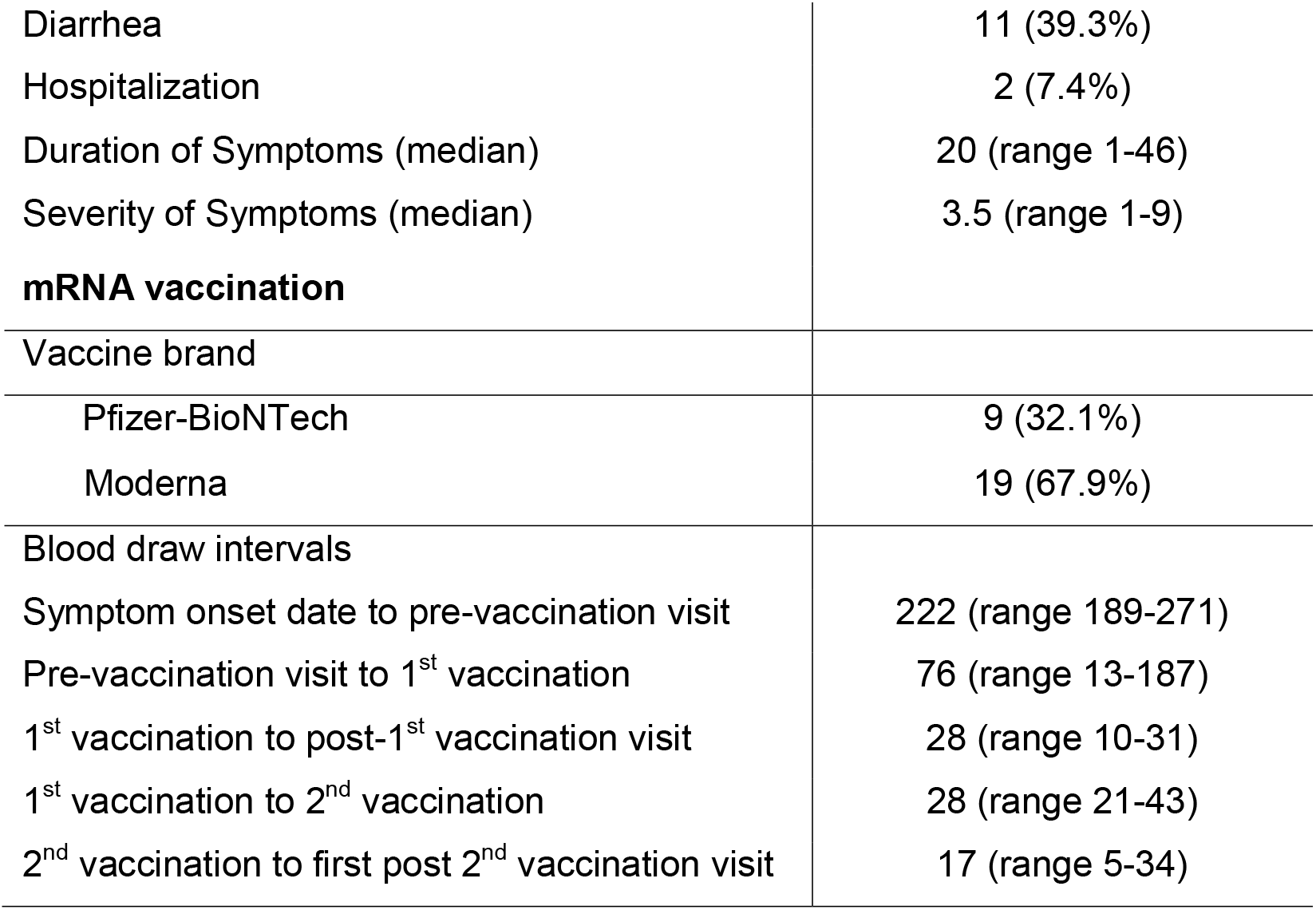
COVID-19 vaccinee characteristics.

**Supplemental table 3:** Naïve vaccinee characteristics.

|  | Naïve vaccinees |
| --- | --- |
|  | n= 18 |
| **Demographics** |  |
| Median Age (years) | 39.8 (range 22-77) |
| Sex |  |
| Men | 11 (61.1%) |
| Women | 7 (38.9%) |
| Race |  |
| White | 11 (55.6%) |
| Black/African American | 0 (0.0%) |
| Asian | 5 (27.8%) |
| Other | 0 (0.0%) |
| Prefer not to say | 2 (11.1%) |
| Hispanic or Latino | 4 (22.2%) |
| **General Characteristics** |  |
| Number of Household Members (median) | 4 (range 2-6) |
| Healthcare worker/in health care field | 12 (66.67%) |
| **Clinical Characteristics** |  |
| Median BMI | 22.1 (range 19.1-32.6) |
| Lung Disease | 1 (5.6%) |
| Diabetes | 1 (5.6%) |
| Cardiovascular Disease | 1 (5.6%) |
| Immunocompromised | 0 (0.00%) |
| **mRNA vaccination** |  |
| Vaccine brand |  |
| Pfizer-BioNTech | 14 (77.8%) |
| Moderna | 4 (22.2%) |
| Blood draw intervals |  |
| Pre-vaccination visit to 1<sup>st</sup> vaccination | 5 (range -1-266) |
| 1<sup>st</sup> vaccination to post-1<sup>st</sup> vaccination visit | 22 (range 10-31) |
| 1<sup>st</sup> vaccination to 2<sup>nd</sup> vaccination | 21 (range 20-43) |
| 2<sup>nd</sup> vaccination to first post 2<sup>nd</sup> vaccination visit | 28 (range 5-34) |

